# Pre-stimulus Activity Mediates Event-Related Theta Synchronization and Alpha Desynchronization During Memory Formation in Healthy Aging

**DOI:** 10.1101/2024.07.17.603896

**Authors:** Dawid Strzelczyk, Nicolas Langer

**Affiliations:** Methods of Plasticity Research, Department of Psychology, University of Zurich, Zurich, Switzerland; University Research Priority Program (URPP) Dynamics of Healthy Aging, Zurich, Switzerland; Neuroscience Center Zurich (ZNZ), Zurich, Switzerland

## Abstract

The capacity to learn is a key determinant for the quality of life but is known to decline to varying degrees with age. However, despite mounting evidence of memory deficits in older age, the neural mechanisms contributing to successful or impeded memory remain unclear. Previous research has primarily focused on memory formation through remembered versus forgotten comparisons, lacking the ability to capture the incremental nature of learning. Moreover, previous EEG studies have primarily examined oscillatory brain activity during the encoding phase, such as event-related synchronization (ERS) of mid-frontal theta and desynchronisation (ERD) of parietal alpha, while neglecting the potential influence of pre-stimulus activity. To address these limitations, we employed a sequence learning paradigm, where 113 young and 117 older participants learned a fixed sequence of visual locations through repeated observations (6423 sequence repetitions, 55 ’944 stimuli). This paradigm enabled us to investigate mid-frontal theta ERS, parietal alpha ERD, and how they are affected by pre-stimulus activity during the incremental learning process. Behavioral results revealed that young subjects learned significantly faster than older subjects, in line with expected age-related cognitive decline. Successful incremental learning was directly linked to decreases of mid-frontal theta ERS and increases of parietal alpha ERD. Notably, these neurophysiological changes were less pronounced in older individuals, reflecting a slower rate of learning. Importantly, the mediation analysis revealed that in both age groups, mid-frontal pre-stimulus theta partially mediated the relationship between learning and mid-frontal theta ERS. Furthermore, the overall impact of learning on parietal alpha ERD was primarily driven by its positive influence on pre-stimulus alpha activity. Our findings offer new insights into the age-related differences in memory formation and highlight the importance of pre-stimulus activity in explaining post-stimulus responses during learning.

## 1. Introduction

The rising prevalence of physical and cognitive impairments among the aging population is placing a growing burden on society ^1,2^. Given the association between older age and declines in learning and episodic memory, it is crucial to enhance our understanding of the neural processes underlying successful or impeded learning in older individuals. However, despite mounting evidence of memory deficits in older age, the precise mechanisms contributing to successful or impeded learning remain elusive ^3–5^. To investigate the processes underlying memory formation, researchers have widely used the subsequent memory effect (SME) paradigm ^6^. This paradigm compares neural activity during the encoding of trials that are later remembered to those that are forgotten. Studies have demonstrated that oscillatory activity, particularly event-related synchronization of mid-frontal theta (i.e., mid-frontal theta ERS) and desynchronization of parietal alpha (i.e., parietal alpha ERD), contribute to improved memory performance ^6–22^.

Mid-frontal theta ERS has been found to play a crucial role in memory formation, showing increased activity during encoding for items that were successfully remembered compared to those forgotten ^6,7,10,19,20,23,24^. This mid-frontal theta ERS increase is directly linked to the amount of information held in working memory ^25^. Current research suggests that theta oscillations coordinate different regions to support associative memory formation and retrieval ^26^. Instead of directly reflecting memory functions, theta is associated with broader processes such as attention and cognitive control ^23,27^. To ensure adequate allocation of attention and cognitive control, crucial functions for managing and directing the flow of information necessary for memory formation, mid-frontal theta fosters coordination among various brain structures. Specifically, mid-frontal theta integrates different structures in the hippocampal formation, the medial temporal lobe and the neo-cortex into coherent neurocognitive networks and binds new information into coherent memory traces^6,7,10,23,24,27,28^.

Parietal alpha ERD also has been found to play an important role in memory formation, facilitating efficient information processing, with items remembered showing higher parietal alpha ERD during encoding than those forgotten ^12–18,22^. Parietal alpha ERD is thought to reflect a reduction in synchronized activity across neural networks, enhancing signal-to-noise ratio critical for creating detailed memories ^13,16^. Moreover, the magnitude of parietal alpha ERD is directly linked to the depth of elaborative encoding and predictive of memory performance, demonstrating its role in supporting the representation of information within episodic memories. Alpha decreases are not stimulus-specific but serve a general process by inhibiting task irrelevant networks, thereby improving the conditions for information representation across various tasks and modalities ^13,16^. This generalized mechanism is essential for controlling the information flow in cortical groups involved in learning, underpinning the enhanced processing depth and volume of information accessible during encoding and retrieval.

While studies employing the SME paradigm have significantly contributed to our understanding of memory formation, they also present several limitations. Firstly, the SME paradigm lacks the capability to capture the incremental nature of learning, because each stimulus is presented only once during the learning phase. Repetition plays a fundamental role in consolidating and strengthening memories, and the lack of stimulus repetition within the SME paradigm hinders our comprehensive understanding of the incremental learning process. Secondly, the majority of research on SME in aging has employed fMRI, with only a handful of EEG studies exploring age-related differences. Moreover, the few EEG studies exploring age-related differences yield inconclusive results, with some reporting attenuated theta ERS in older age ^29,30^, while others find no significant differences, as evidenced by a range of studies ^11,15,31–33^. Similarly, while some studies have reported a diminished alpha ERD in older individuals ^11,15,34^, other studies have not identified any significant differences in alpha ERD between age groups ^32^. Thirdly, the ERS/ERD are relative measures of brain activity and reflect a combination of pre- and post-stimulus activity. Consequently, it is difficult to determine whether the relationship between ERS/ERD and behavior is driven by pre-stimulus activity, post-stimulus activity or both. This aspect is especially relevant in studies of repetition learning, including the present study, because the pre-stimulus activity may represent accumulated knowledge or expectations about upcoming stimuli. Subtracting baseline activity from post-stimulus activity (i.e., ERD/ERS) overlooks the influence of pre-stimulus neural activity on the incremental learning process. However, whether pre-stimulus activity reflects general memory-promoting state or it is specifically linked to cognitive processes that are related to learning and memory formation, is unclear^10^.

Thus, we recruited a large sample of 113 young and 117 older participants and employed a sequence learning paradigm that allowed us to investigate the modulation of pre-stimulus activity over the course of learning and its effects on post-stimulus ERS and ERD responses. This approach provides valuable insights into the anticipatory and preparatory cognitive processes occurring before stimulus presentation. Furthermore, in response to previous studies that reported age-related flattening of the 1/f slopes and a slowing of of the alpha rhythm ^35,36^, we disentangled the EEG signal into periodic and aperiodic components and extracted individual alpha frequencies (IAF) for all subsequent analyses. Given the role of mid-frontal theta ERS in memory formation, we hypothesized that mid-frontal theta ERS decreases over the course of learning. The difficulties of older subjects with binding the items into coherent memory traces will be reflected by lower mid-frontal theta ERS and more gradual decrease over the course of learning. Given that parietal alpha ERD promotes the depth of information processing and determines the amount of information available at encoding and retrieval, we further hypothesize that the parietal alpha ERD to be high at the beginning of learning, and gradually decrease (i.e., reducing desynchronisation), as the amount of information to process decreases (i.e., memory strengthens). We also expected parietal alpha ERD to be attenuated in the older group reflecting less efficient information processing. Finally, to determine whether the relationship between ERS/ERD and behavior is contributed by pre-stimulus activity, post-stimulus activity or both, we performed causal mediation analysis to investigate whether the effect of learning progress on mid-frontal theta ERS and parietal alpha ERD is mediated by pre-stimulus activity.

## 2. Methods

### 2.1. Participants

In the present study 113 young (age range 19 - 43 years; mean age, 24.63 ± 4.52 years, 45 male, 88 right-handed) and 117 older (58 - 84 years; mean age, 69.1 ± 5.31 years, 53 male, 95 right-handed) participants were recruited. Table 1 shows basic demographic details for both age groups. All participants were healthy, reported normal or corrected to normal vision and no current neurological or psychiatric diagnosis. The young group consisted of graduate students at the University of Zürich or other universities nearby. The older subjects were recruited through announcements in newsletters and during lectures within the Senior-University of Zürich. As a compensation the participants were given course credit or monetary reward (25 CHF/h). This study was conducted according to the principles expressed in the Declaration of Helsinki. The study was approved by the Institutional Review Board of Canton Zurich (BASEC - Nr. 2017 - 00226). All participants gave their written informed consent before participation.

**Table 1.**
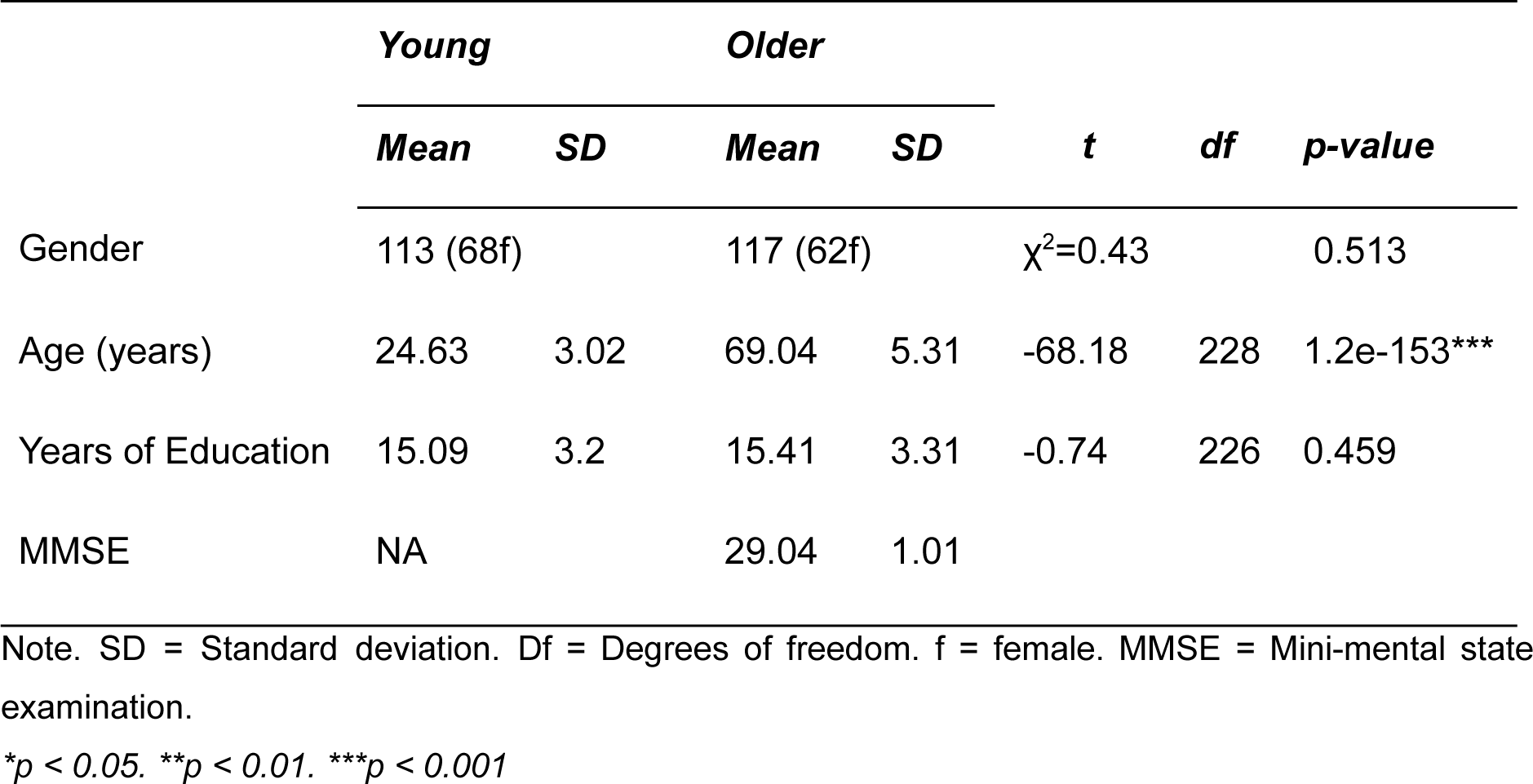
Demographics information.

|  | *Young* |  | *Older* |  | *t* | *df* | *p-value* |
| --- | --- | --- | --- | --- | --- | --- | --- |
|  | *Mean* | *SD* | *Mean* | *SD* |  |  |  |
| Gender | 113 (68f) | | 117 (62f) | | $\chi^2=0.43$ | | 0.513 |
| Age (years) | 24.63 | 3.02 | 69.04 | 5.31 | -68.18 | 228 | 1.2e-153*** |
| Years of Education | 15.09 | 3.2 | 15.41 | 3.31 | -0.74 | 226 | 0.459 |
| MMSE | NA |  | 29.04 | 1.01 |  |  |  |

Note. SD = Standard deviation. Df = Degrees of freedom. f = female. MMSE = Mini-mental state examination.
\* $p < 0.05$ . \*\* $p < 0.01$ . \*\*\* $p < 0.001$

### 2.2. Procedure

The data used in this study was recorded in our laboratory in the context of a larger project that aims to quantify age effects on eye movement behavior and electroencephalography (EEG) recordings of resting-state and other task-based paradigms ^37,38^. The data was collected in two experimental sessions separated by a week. Upon arrival, the older participants performed the Mini-mental State Exam (MMSE) in order to screen for cognitive impairment and dementia ^39^. All participants accomplished a MMSE score above the threshold of 25.

During the EEG acquisition, the participants were comfortably seated in a chair in a sound- and electrically shielded Faraday recording cage. The cage was equipped with a chinrest to minimize head movements and a 24-inch monitor (ASUS ROG, Swift PG248Q, display dimensions 531 x 299 mm, resolution 800 x 600 pixels resulting in a display: 400 x 298.9 mm, vertical refresh rate of 100 Hz) on which the experiment was presented. The distance between the chinrest and the monitor was 68 cm.

### 2.3. Sequence Learning Task

A visual sequence learning paradigm was first developed by Moisello et al. (2013) and is currently considered as an important tool in assessing reliable indices of memory formation and learning progress ^37,41^. Through the employment of simplified stimuli, this paradigm has the advantage of mitigating confounding effects of stimuli properties such as memorability, semantic content, and sensory characteristics that typically influence learning efficacy. By enabling within-sequence comparisons among standardized stimuli, it ensures a targeted investigation of the neural mechanisms underpinning learning success ^42,43^. The participants were asked to learn a fixed sequence of eight visual stimulus positions (Figure 1A). The stimuli consisted of filled white circles (visual angle of 0.84°) and were presented on a computer screen with a bright gray background, positioned equidistant around a ring of fixed eccentricity (visual angle of the distance between center of the screen and the stimulus of 4.21°). Each stimulus was presented for 600 ms with an offset-to-onset interval of 1300 ms. Before the main task recording, a training task was administered, consisting of 4 stimuli placed on the same 8 locations, in order to familiarize the participants with the tasks and to ensure task comprehension. The participants performed the training task until they correctly recalled all 4 locations. Feedback was provided only during the training task.

**Figure 1:**
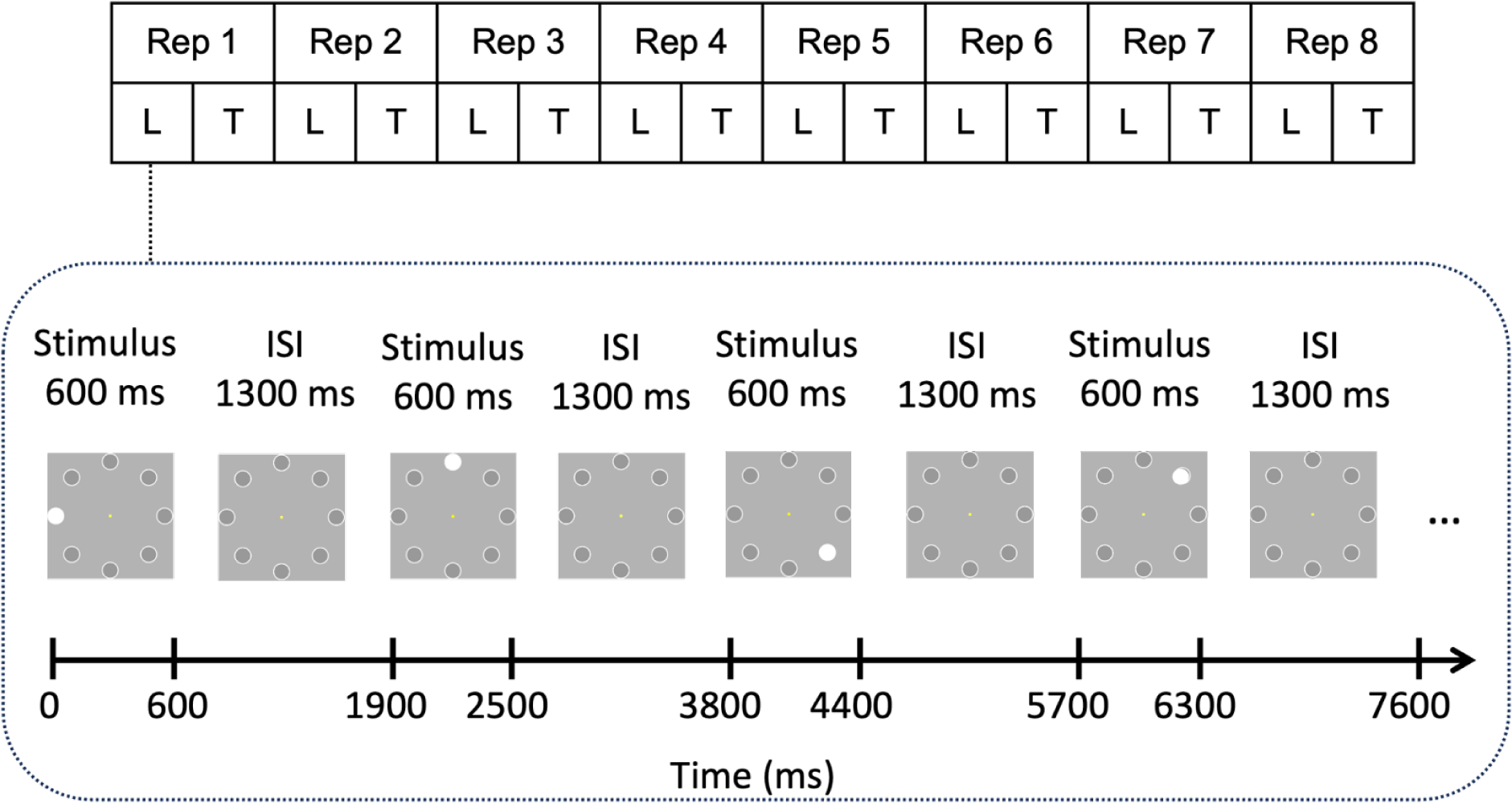
Sequence learning task and the design of the present study. (A) A sequence consisted of eight positions at which white circles were presented one by one. Each stimulus was presented for 600 ms with the interstimulus interval of 1300 ms. (B) Each sequence repetition (Rep) consisted of a learning phase (L) in which all sequence elements were presented and a testing phase (T) in which the participants were asked to recall the position of the stimuli. Adapted from Strzelczyk et al. (2023) licensed under CC BY 4.0.

The main task consisted either of eight sequence repetitions or ended after the participant correctly recalled the sequence of stimuli three times in a row. Each sequence repetition consisted of a learning phase and a testing phase. In the learning phase, the participants were told to focus on a yellow dot at the center of the screen (controlled by an eye-tracking device) and memorize the position of each stimulus. In the test phase, the participants attempted to recall the sequence using a computer mouse by clicking the locations on a computer screen. There was no time restriction on providing responses, and no feedback on their performance was given. Therefore, the duration for learning a sequence varied between 2 and 5 min, depending on the speed of recall reports and number of sequence repetitions (i.e., 3 to 8 repetitions). Overall each participant learned six different sequences, resulting in a minimum of 18 (in case the participant solves everything correctly from the beginning) and maximum of 48 sequence repetitions.

### 2.4. Behavioral Data

To quantify learning performance over sequence repetitions, *accuracy* was computed for each participant. The *accuracy*, which reflects the cumulative sequence knowledge, was defined as the ratio of the number of correct nCorrect(P, sr) responses to the total number of stimuli nT(P, sr) in each sequence repetition, where *P* represents the participant and *sr* the sequence repetition.

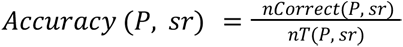

The participants were not instructed to prioritize speed in their responses. Consequently, the reaction time data should be interpreted with caution. Due to this consideration, we have chosen to include the results of the reaction time analysis in the supplementary materials (Supplementary Figure 1).

### 2.5.1. Eye-Tracking data acquisition

An infrared video-based eye tracker (EyeLink 1000 Plus, SR Research; http://www.sr-research.com/) recorded eye movements at a sampling rate of 500 Hz and an instrumental spatial resolution of 0.01°. The eye tracker was calibrated and validated before each task with a 9-point grid until the average error for all 9 points was below 1°. In this study eye tracker data were used in a control analysis accounting for whether the participants maintained focus at the center of the screen as instructed (i.e., yellow dot). Therefore, eye movements during each stimulus presentation period were included into the models as a covariate (i.e., 1 = kept fixation, 0 = lost fixation). It was deemed that a participant had kept fixation, if the gaze was directed at the center of the screen within a square of side 1.26° of visual angle during at least 90% of the total duration of the stimulus presentation period.

#### 2. 5. 2. EEG data acquisition and preprocessing

EEG data were recorded at a sampling rate of 500 Hz using a 128-channel Hydrogel net system (Electrical Geodesics Inc.). The recording reference was at Cz (vertex of the head), and impedances were kept below 40 kΩ. All data were analyzed using Matlab 2020a (The MathWorks, Inc., Natick, Massachusetts, United States) and RStudio 4.0.2 (R Core Team). Due to technical difficulties (EEG data not saved properly, missing subjects’s responses, Matlab crash during experiment, subject attending only 1 session (i.e., 18 subjects)) data from 118 sequences was not available, resulting in a total of 1262 sequences (i.e., 6423 sequence repetitions, 55 ’944 stimuli). The data were preprocessed in Automagic 2.6.1, a MATLAB based toolbox for automated, reliable and objective preprocessing of EEG-datasets^44^. In the first step in Automagic, the bad channels were detected using the PREP pipeline ^45^. A channel was defined as bad based on 1) extreme amplitudes (z-score cutoff for robust channel deviation of more than 5), 2) lack of correlation (at least 0.4) with other channels with a window size of 1s to compute the correlation, 3) lack of predictability by other channels (channel is bad if the prediction falls below the absolute correlation of 0.75 in a fraction of 0.4 windows of a duration of 5s), 4) unusual high frequency noise using a z-score cutoff for SNR of 5. These channels were removed from the original EEG data. The data was filtered using a high-pass filter with 0.5 Hz cutoff using the EEGLAB function pop_eegfiltnew ^46^. Line noise was removed using a ZapLine method with a passband edge of 50 Hz ^47^, removing 7 power line components. Next, independent component analysis (ICA) was performed. However, as the ICA is biased towards high amplitude and low frequency noise (i.e., sweating), the data was temporarily filtered with a high-pass filter of 1 Hz in order to improve the ICA decomposition. Using the pre-trained classifier IClabel ^48^ each independent component with a probability rating >0.8 of being an artifact such as muscle activity, heart artifacts, eye activity, line noise and channel noise were removed from the data. The remaining components were back-projected on the original 0.5 Hz high-pass filtered data. In the next step, the channels identified as bad were interpolated using the spherical interpolation method. Finally, the quality of the data was automatically and objectively assessed in Automagic, thus increasing research reproducibility by having objective measures for data quality. Using a selection of 4 quality measures the data was classified into three categories: Good (1091 sequences), OK (146 sequences) or Bad (25 sequences). Data was classified as bad, if 1) the proportion of high-amplitude data points (>30μV) in the signal is greater than 0.3, or 2) more than 30% of time points show a variance greater than 30μV across all channels, or 3) 30% of the channels show variance greater than 30μV, or 4) the ratio of bad channels is greater than 0.3. For the further analysis only the datasets with Good and OK ratings were used.

Subsequently, 23 channels were excluded from further analysis, including 10 EOG channels and 13 channels located on the chin and neck as they capture little brain activity and are mostly contaminated with muscle artifacts ^35,42,49^. Next, the data was re-referenced to average reference and segmented from -750 to 1150 ms after stimulus onset (i.e., presentation of white circle on one of the eight positions). The segments were inspected using an amplitude threshold of 90μV resulting in an average rejection rate of 10.7% per participant.

#### 2. 5. 3. Spectral analysis

The spectral analysis was conducted using the MATLAB FieldTrip toolbox ^50^. For each sequence repetition (i.e., 8 trials), spectral analysis was computed using the fast Fourier transform (FFT) algorithm, employing a sliding window of 500 ms and applying a Hanning taper to achieve a frequency smoothing of approximately 3 Hz. Power estimates were calculated within a latency window ranging from -750 to 1150 ms after stimulus onset, with steps of 50 ms, and then averaged across repetitions. The segment length was carefully selected to align with the experimental design and analysis requirements: 1) The stimulus-to-stimulus interval was at 1900 ms, 2) a 500 ms sliding window for FFT analysis was employed. An important aspect to highlight is the effective range of the power estimates obtained. Due to the nature of the sliding window approach in FFT computation, the first 250 ms of the segment (from -750 ms to -500 ms) are utilized for the initial FFT calculation. As a result, the actual range of power estimates effectively starts only from -500 ms. Therefore, the true range of obtained power estimates spans from -500 to 900 ms post-stimulus. The estimated frequency range spanned from 1 to 40 Hz in steps of 2 Hz. Zero padding was applied to provide a frequency resolution of 0.2 Hz. Following this, the power estimates were further decomposed into periodic and aperiodic components using the SpecParam algorithm implemented in FieldTrip ^36^. This step is crucial, when comparing different age groups, given that older individuals tend to display flatter 1/f slopes than younger counterparts ^35,51^. Therefore, the main emphasis of our study was placed on the aperiodic adjusted signal. The non-adjusted signal, by contrast, was utilized solely for control analysis purposes.

#### 2. 5. 4. SpecParam algorithm and aperiodic adjusted power

The SpecParam algorithm ^36^ was employed to parameterize the power spectrum, effectively separating oscillatory components from the aperiodic signal. This algorithm estimates oscillatory peaks that are overlaid on the aperiodic signal, and thus, these peaks are measured relative to the aperiodic signal rather than absolute zero. The parameterization of the Power Spectral Density (PSD) is achieved through an iterative process, wherein the algorithm fits the aperiodic signal represented by *L* to the observed smoothed spectral signal, resulting in two parameters: the aperiodic intercept *b* and the aperiodic exponent *x* (i.e., negative slope, with smaller *x* indicating a flatter spectrum). Mathematically, it can be expressed as:

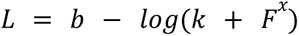

Here, *F* represents the vector of input frequencies, and *k* is the “knee” parameter, which, in this context, is set to 0 in the proposed analysis. This means that no bend of the aperiodic component is additionally modeled in the data, adhering to the default state of the SpecParam algorithm.

To extract oscillatory components, this aperiodic signal is subtracted from the power spectrum. Gaussians are then iteratively fitted to the remaining signal, and they are subtracted whenever data points exceed two standard deviations of the data. These fitted Gaussians represent the true oscillatory components in the data, while data points below the specified threshold are considered as noise. Consequently, this process yields a data-driven number of Gaussians, each characterized by the frequency center, power relative to the aperiodic signal, and the frequency bandwidth. The power spectrum is, therefore, modeled as follows:

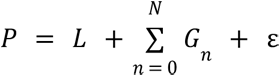

Here, *G_n_* represents the *n_th_* Gaussian, and *ε* accounts for the noise not captured by the model. It’s worth noting that this description of the algorithm is a simplification, and for a more detailed definition, you can refer to the original publication ^36^. The specParam algorithm was applied within a frequency range of 1 to 40 Hz, chosen to avoid overfitting of noise as small bandwidth peaks in very low frequencies. The algorithm settings included: peak width limits set to [1, 8]; no limit on the maximum number of peaks; a minimum peak height set at 0; a peak threshold defined as 2 standard deviations above the mean; and the aperiodic mode set to ‘fixed’.

Finally, to adjust the power measurements for the aperiodic signal, the aperiodic component was first reconstructed using its parameters. This reconstructed aperiodic signal was then subtracted from the total power spectrum, yielding an aperiodic-adjusted power spectrum. The aperiodic adjusted power spectrum then was used for further analysis. The extracted aperiodic component was not the focus of this work and was only briefly described in the supplementary material (Supplementary Material 5.7).

Due to the sliding window approach, spectral analysis yielded specParam model fits (i.e., R^2^) ranging from -500 to 900 ms around stimulus onset, in 50 ms intervals, for each channel and sequence repetition. To compute the average specParam model fit for each sequence repetition, we averaged the fits across the time points and channels, resulting in one specParam model fit per sequence repetition. Out of 6423 sequence repetitions across all subjects, 391 (6.1%) showed an average specParam model fit below 0.9 and were excluded from further analysis (39 repetitions in young subjects and 352 repetitions in older subjects). The remaining sequence repetitions demonstrated a high overall specParam model fit, both in young (M = 0.9464, SD = 0.0127) and older (M = 0.9355, SD = 0.0157) subjects.

#### 2. 5. 5. Channel and time window selection

Having isolated the periodic and aperiodic components, our next objective was to identify channels and time windows for further statistical analysis. To achieve this, we first computed the grand average aperiodic-adjusted power spectrum over all subjects, repetitions and channels. We observed a pronounced, ongoing power in the canonical alpha range (i.e., 8 - 13 Hz), marked by a decrease in power starting around 200 ms post-stimulus, and a noticeable activity in canonical theta range (i.e., 4 - 7 Hz) peaking around 250 ms post-stimulus (Supplementary Figure 2). Taking into account that, based on the grand average, the decrease in alpha power began 200 ms post-stimulus, and theta peaked at 250 ms, and given that the sliding window for FFT was 500 ms, with the epoch extending from -500 ms to 900 ms post-stimulus, we selected the time window for parietal alpha analysis from 250 to 900 ms post-stimulus and for mid-frontal theta analysis from 100 to 500 ms post-stimulus. Next, we plotted topographical maps of parietal alpha in the time window of 250 - 900 ms and mid-frontal theta in the time window of 100 - 500 ms to identify channels for further analysis. To account for inter-individual anatomical differences and enhance the robustness of the electrode cluster, as well as to address the altered topography of alpha due to the circular presentation of stimuli ^52,53^, we identified the following channels for further analysis of parietal alpha: E75, E70, E83, E62, E72, E67, E71, E77, E76, E52, E65, E60, E61, E59, E66, E78, E90, E92, E85, E84, E91, E58, E96. Next, we identified the following channels for further analysis of mid-frontal theta: E5, E6, E7, E11, E12, E106, E13, E112 (see supplementary Figure 2 for the electrode positions of parietal alpha and mid-frontal theta). The identified time windows and channels resonate with those frequently employed in prior studies ^9,23^, further validating our methodological choices and ensuring that our findings are comparable and relevant within the broader research context.

#### 2. 5. 6. Computation of individual alpha peak frequency

Subsequently, we proceeded with the computation of the individual alpha peak frequency (IAF). IAF measure was selected instead of a canonically defined alpha range (i.e., 8 - 13 Hz), as the shift of the IAF during aging might introduce a bias when power is averaged within a fixed-frequency window ^35,54^. Specifically, it has been shown that the IAF decreases as individuals age and the rate of decline can vary among individuals. To accurately estimate IAF, high frequency resolution is required. While padding a signal with zeros increases the number of data points and smooths the power spectrum, the added zeros do not contain any real information about the signal. Therefore, to truly obtain high frequency resolution, we concatenated all trials from session 1 and session 2 and treated the data as a single trial. Please note that this analysis was only performed to estimate IAF. All subsequent statistical analyses (i.e., mid-frontal theta and parietal alpha over sequence repetitions and mediation analyses) were performed on the zero-padded data described in section 2. 5. 3. To estimate IAF, spectral analysis was computed using the fast Fourier transform algorithm multiplied by multitapers (i.e., DPSS) resulting in frequency resolution of 0.2 Hz. Following this, the power estimates were further decomposed into periodic and aperiodic components using the specParam algorithm implemented in FieldTrip ^36^. The decomposition showed very high specParam model fits, both in young (M = 0.9921, SD = 0.0014) and older subjects (M = 0.9852, SD = 0.0035). The IAF was found by determining the frequency of maximum power between a lower and upper frequency limit. Following previous work, these frequency limits were set to 7 and 14 Hz ^8,55^. If the peak was located outside of the preselected frequency range or at the border, no alpha peak was extracted for that subject, and the corresponding data was excluded from further analysis. In total, IAF was found for 199 subjects, comprising 105 young (i.e., 93.75% detection rate) and 94 older (i.e., 81.03% detection rate) participants. Subjects without a detectable IAF were excluded from further analyses. The average IAF was 10.5277 ± 1.2478 Hz for the young group and 10.2362 ± 1.4032 Hz for the older group (M ± SD). The difference in IAF between age groups did not reach significance (t = 1.5512; p = 0.1225; CI = [-0.0791, 0.6621]).

#### 2. 5. 7. Extraction of power for statistical analyses

Event-related synchronization and desynchronization were computed by comparing the power in the frequency band of interest during the event-related period to the power during the baseline period. In the present study, we initially adjusted the baseline within each repetition, aiming to replicate previous findings related to the subsequent memory effect (SME) on ERS/ERD. To achieve this, we focused solely on the event-related activity and corrected the baseline within each sequence repetition (i.e., adjusted the pre-stimulus activity of each sequence repetition to 0). As the output of the specParam algorithm in FieldTrip was already on the logarithmic scale (dB), the formula for calculating ERS/ERD on aperiodically adjusted power was as follows:

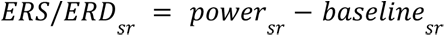

where *sr* represents the sequence repetition. As part of a sensitivity analysis, we also investigated the conventional power without adjusting for the aperiodic component (Supplementary Material 5. 3). Since this power was not yet on a logarithmic scale, we first corrected the baseline of the data and then converted it to a logarithmic scale as follows:

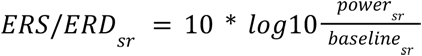

Next, to investigate changes in the pre-stimulus aperiodic adjusted activity over the course of learning and its effects on post-stimulus ERS/ERD, we corrected the baseline within each subject and allowed the pre-stimulus activity to vary within the subject across sequence repetitions. We first computed the *average baseline* across all sequence repetitions of a stimulus sequence and subtracted this value from each sequence repetition. Subsequently, we extracted the baseline values and used them in causal mediation analysis to investigate whether the baseline affects ERS/ERD.

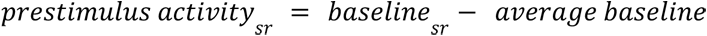

The chosen baseline interval ranged from -500 ms to -250 ms prior to stimulus onset. This baseline interval was specifically chosen to align with our Fast Fourier Transform (FFT) analysis and to avoid overlap with the post-stimulus activity, ensuring a clear delineation between pre-stimulus baseline and stimulus-evoked responses. Next, we proceeded with the extraction of power estimates for further statistical analysis. The power was individually extracted based on each subject’s IAF. Specifically, parietal alpha power was extracted by averaging the power within the frequency window ranging from -4 Hz to +2 Hz, relative to the subject’s IAF, and within the time window of 250 to 900 ms ^8^. Similarly, mid-frontal theta power was extracted by averaging the power within the frequency window from -6 Hz to -4 Hz, relative to the IAF, and within the time window of 100 to 500 ms ^8^.

#### 2. 5. 8. Simulation study to explore the impact of baseline correction on event-related synchronization and desynchronization

As outlined in the introduction, event-related synchronization and desynchronization (ERS/ERD) are relative measures of brain activity and reflect a combination of pre- and post-stimulus activity. Consequently, it is difficult to determine whether the relationship between ERS/ERD and behavior is contributed by pre-stimulus activity, post-stimulus activity or both. This consideration is particularly relevant in our study, where the pre-stimulus activity might reflect the individual’s knowledge or expectations about next stimuli. To shed light on this issue, we performed a simulation study, which is depicted in Figure 2. Panel A illustrates the typical temporal evolution of event-related desynchronization over time in the first repetition (represented by a thick black line) alongside four hypothetical scenarios depicting how power might evolve in the second repetition (indicated by dashed lines). Specifically, these scenarios include an increase in power pre-stimulus (scenario A), a decrease post-stimulus (scenario B), an increase pre-stimulus followed by a decrease post-stimulus (scenario C), and a decrease in both pre- and post-stimulus phases (scenario D). Panel B illustrates the state-of-the-art ERD approach, where a baseline correction is performed within each repetition (i.e., condition), where pre-stimulus activity is adjusted to 0.

**Figure 2:**
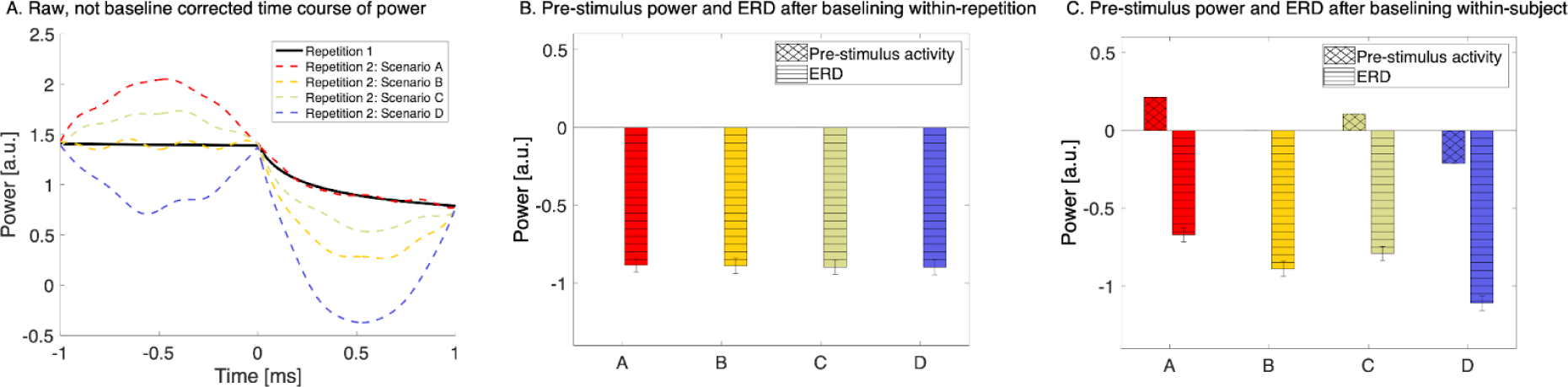
Simulation study illustrating the impact of baseline correction on event-related activity. Panel A displays the typical temporal progression of event-related desynchronization (ERD) in the first repetition (shown as a thick black line) and contrasts it with four hypothetical scenarios for power evolution in the second repetition (depicted by dashed lines). These scenarios encompass: an increase in power before the stimulus (scenario A), a decrease after the stimulus (scenario B), an increase before and decrease after the stimulus (scenario C), and a decrease in both pre- and post-stimulus phases (scenario D). Panel B shows baseline correction applied within each repetition, adjusting pre-stimulus activity to zero, highlighting that such a correction alone does not conclusively indicate whether power changes occurred pre- or post-stimulus, as all ERDs appear similar. Panel C demonstrates the application of baseline correction within each subject, enabling the distinction of whether power changes occur before or after the stimulus.

It is important to note that solely by examining the event-related desynchronisation (as seen in Panel B), we cannot definitively determine whether the observed power change occurred pre- or post-stimulus since all ERDs appear identical. However, when we apply a baseline correction within each subject (Panel C), we gain the ability to discern whether the power change occurred pre- or post-stimulus. In essence, this simulation study underscores the critical role of baseline correction in dissecting the temporal dynamics of ERS/ERD and understanding whether changes in brain activity are driven by pre-stimulus or post-stimulus processes.

### 2. 6. Statistical analysis

For each statistical analysis, we first identified the best generalized linear mixed-effects model for explaining the data. In the following formulas fixed effects are denoted by a “+” symbol and interaction effects by an “∗” symbol, in line with the Wilkinson notation ^56^. The models were as general as possible at first and were progressively simplified using the lmerTest::step function for linear mixed effect models in R Studio ^57^ in order to identify the best-fit model. The lmerTest::step function works by iteratively removing one variable from the model and fitting a reduced model. Then, an F-test is performed between the full model and the reduced model, and the p-value is calculated using Satterthwaite’s approximation. If the p-value is below significance level (0.05), the variable is deemed to be important for the model and is retained. If the p-value is above the significance level, the variable is deemed to be non-significant and is removed from the model. This process is repeated for each variable in the model until no further variables can be removed ^57^. The predictors included the repetition number (continuous variable: 1-8), age group (factor of 2 levels: young, older), pre-stimulus power (continuous variable), session (factor of 2 levels: 1, 2), eye movements (factor of 2 levels: kept, lost fixation), subject (factor of 199 levels), sequence number (factor of 6 levels: 1-6, i.e., each participant attempted to memorize six sequences). The fixed and random effects of the model were specified depending on the goal of the analysis, keeping in mind that a mixed model requires at least 5 levels for a random intercept term to accurately estimate the variance ^58^.

#### 2. 6. 1. Behavioral analysis

We began the behavioral analysis by examining the learning progress of both age groups. Learning performance was formally modeled as the cumulative knowledge about the sequence (i.e., accuracy). We specified a linear mixed effect model with an accuracy as dependent variable (continuous variable: 0-1). The fixed effects included the sequence repetition number, age group, interaction of sequence repetition number and age group, session, gender and the random effects included subject and sequence number.

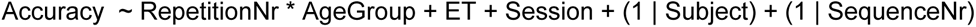

#### 2. 6. 2. EEG data analysis

##### Mid-frontal theta ERS and parietal alpha ERD over sequence repetitions

We proceeded with the analysis of EEG data by investigating the changes of mid-frontal theta ERS and parietal alpha ERD over the course of learning (i.e., sequence repetitions). We specified two independent linear mixed effect models with a mid-frontal theta ERS and parietal alpha ERD as dependent variables (continuous variables). The fixed effects included the sequence repetition number, age group, interaction of sequence repetition number and age group, session, gender and the random effects included subject and sequence number.

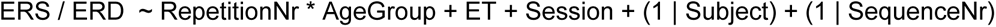

To enhance the robustness of our results, we performed a series of control and sensitivity analysis. First, we replicated the original analysis using conventional power measurements without adjusting for aperiodic components (Supplementary Material 5. 2). Subsequently, we investigated whether the observed changes in mid-frontal theta ERS and parietal alpha ERD across repetitions were directly related to learning or merely a result of habituation and have changed as a function of the time spent on a task. To provide stronger evidence for the link between mid-frontal theta ERS, parietal alpha ERD, and learning processes, we examined if the modulations of these components change as a function of learning state, that is, from stimulus being unknown, to newly learned, known and finally very well known (Supplementary Material 5. 4), independent of sequence repetition ^37,41^. To further substantiate the connection between mid-frontal theta ERS and the process of learning, as well as parietal alpha ERD and learning, we aligned the sequence repetitions to the point where the sequence was correctly recalled for the first time (i.e., considering the number of repetitions before or after this point). The first correctly recalled sequence was labeled as distance 0, with previous repetitions assigned negative distances and subsequent ones positive distances. This approach allowed for a more detailed investigation of the learning process over time, offering finer temporal resolution than our earlier learning state categorizations. For detailed insights, please refer to the supplementary material (Supplementary Material 5. 4 & 5. 5).

##### Mediating effects of pre-stimulus activity on parietal theta ERS and parietal alpha ERD

ERS/ERD represents a combination of pre- and post-stimulus activity. It is unclear whether changes in ERS/ERD originate from pre- or post-stimulus activities. To disentangle these components and assess whether pre-stimulus activity influences the strength of ERS/ERD, we extracted the pre-stimulus activity and used a mediation analysis ^59^ to investigate whether the effect of learning progress (i.e., RepetitionNr) on mid-frontal theta ERS and parietal alpha ERD can be explained by a mediating effect of pre-stimulus power. The mediation analysis was computed for young and older subjects separately to allow for a detailed understanding of the mediation process within each age group. For an analysis that includes age group as a moderating factor (i.e. moderated mediation), please see the supplementary material. To allow interpretation of the estimates and strength comparisons of the effects of each independent variable on dependent variables, we scaled all variables before computing the mediation analysis. First, we defined the M0 model to assess the direct effect of sequence repetition on ERS/ERD disregarding the effect of pre-stimulus power.

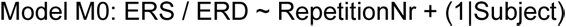

Model M to assess the effect of repetition number on pre-stimulus power was defined as:

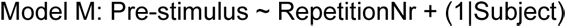

Model Y, which comprises the effect of pre-stimulus power and the effect of repetition number on ERS/ERD, was defined as:

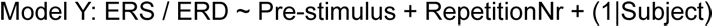

Models M and Y were used as input for mediation analysis. Figure 3 depicts the mediation relation analyzed in this study. The mediation analysis was calculated using the R package for mediation analysis with nonparametric bootstrap for confidence intervals and 1000 Monte Carlo draws ^59^. Prior to the mediation analysis, we confirmed a main effect of repetition number in either Model M0 or M, which is a premise for consecutive mediation analysis ^60,61^. In particular, the main effect of repetition number on pre-stimulus power shows how increasing repetition number is associated with an increased pre-stimulus power.

**Figure 3:**
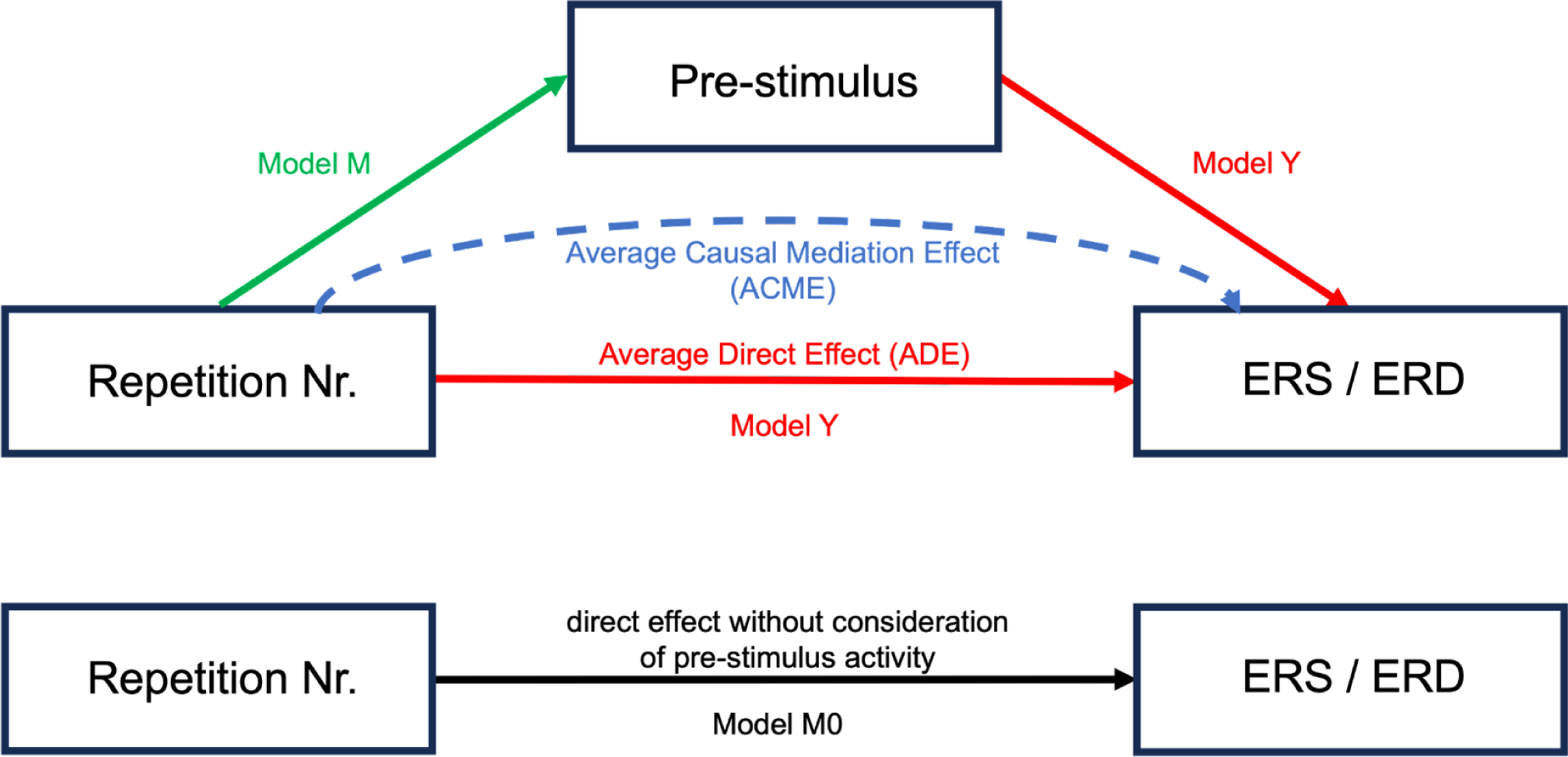
Model Y (red) captures the average direct effect (ADE) of repetition number on ERS/ERD. Model M (green) reflects the effect of repetition number on pre-stimulus power regardless of ERS/ERD. The indirect effect of repetition number on ERS/ERD through the mediation of pre-stimulus power is the average causal mediation effect (ACME; blue). Model M0 (black) captures the relationship of repetition number on ERS/ERD disregarding any effect of pre-stimulus power.

## 3. Results

### 3.1. Behavioral results

Table 1 shows basic demographic information for young and older participants.

#### Visual sequence learning task performance

Across both sessions (i.e., 6 sequences), young subjects required on average 4.45 ± 0.97 repetitions, while older subjects required 6.43 ± 1.10 repetitions to finish the task (M ± SD). The difference was statistically significant (t = -14.27; p = 2.94e-33; CI = [-2.25, -1.70]). The task was considered as finished after 8 repetitions or after the subjects correctly recalled all sequence elements 3 times in a row (i.e., fastest learners completed the task after 3 and slowest learners after 8 repetitions).

We next tested the hypothesis that the knowledge about sequence elements (i.e., accuracy) increases with each sequence repetition and tested for potential age differences. Figure 4 shows the progression of accuracy over the sequence repetition. To visualize the interindividual variance in performance, we further subdivided the participants into groups based on the number of sequence repetitions required for memorization of all sequence elements. The black lines represent the learning curves of each of those groups.

**Figure 4:**
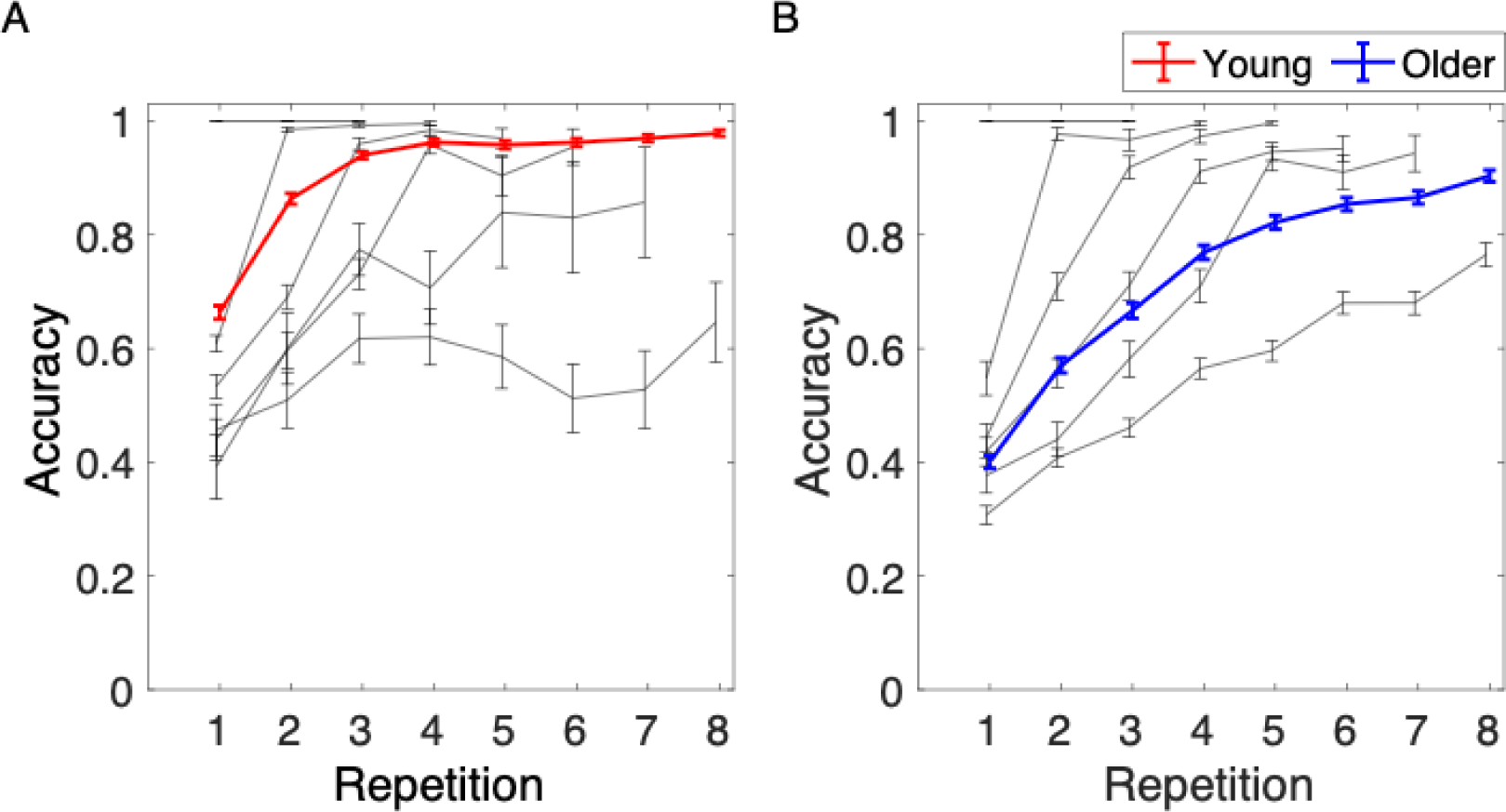
Sequence learning task performance across sequence repetitions in young and older participants. The task lasted 3 to 8 repetitions, depending on learning rate. The black lines indicate the learning curves of participants that completed the task after 3, 4, 5, 6, 7 or 8 sequence repetitions. Error bars represent the standard error of the mean. Adapted from Strzelczyk et al. (2023) licensed under CC BY 4.0.

First, we established the best-fit model for accuracy. The model included fixed effects of repetition number, age group, and their interaction, main effect of eye tracker, and a random effect of subject.

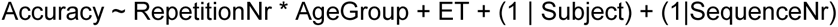

The model revealed a significant main effect of repetition number (β = 0.08; p = 2.4e-94; CI = [0.07, 0.09]), that is the accuracy increased with each sequence repetition (Figure 2A-B). Moreover, there was a significant main effect of age group (β = -0.20 ; p = 1e-18; CI = [-0.25, -0.16]), suggesting a lower accuracy in the older group. There was also a significant interaction of repetition number and age group, indicating a greater increase in accuracy with each sequence repetition in the older group compared to the increase in the young group (β = -0.01; p = 0.001; CI = [-0.02, -0.01]). Finally, there was a significant main effect of eye movements (β = 0.003; p = 0.023; CI = [0.00, 0.01]), that is the accuracy was slightly higher when subjects were fixating the center of the screen. In addition, we observed a substantial variation of accuracy between subjects (SD = 0.12) and sequences (SD = 0.01). Summarized, young and older subjects gradually learn the stimuli over repeated sequence presentations, with the young learning on average faster than older participants.

### 3.2. Mid-frontal theta ERS and parietal alpha ERD over sequence repetitions

In the next step, we tested the hypothesis that the learning progress is accompanied by decreasing mid-frontal theta ERS (i.e., reducing synchronization) and decreasing parietal alpha ERD (i.e., reducing desynchronization). The average time-frequency decompositions in each sequence repetition are presented on Figure 5. Top panel shows the time-frequency decompositions across sequence repetitions in young subjects, and bottom panel shows the time-frequency decompositions in older subjects.

**Figure 5:**
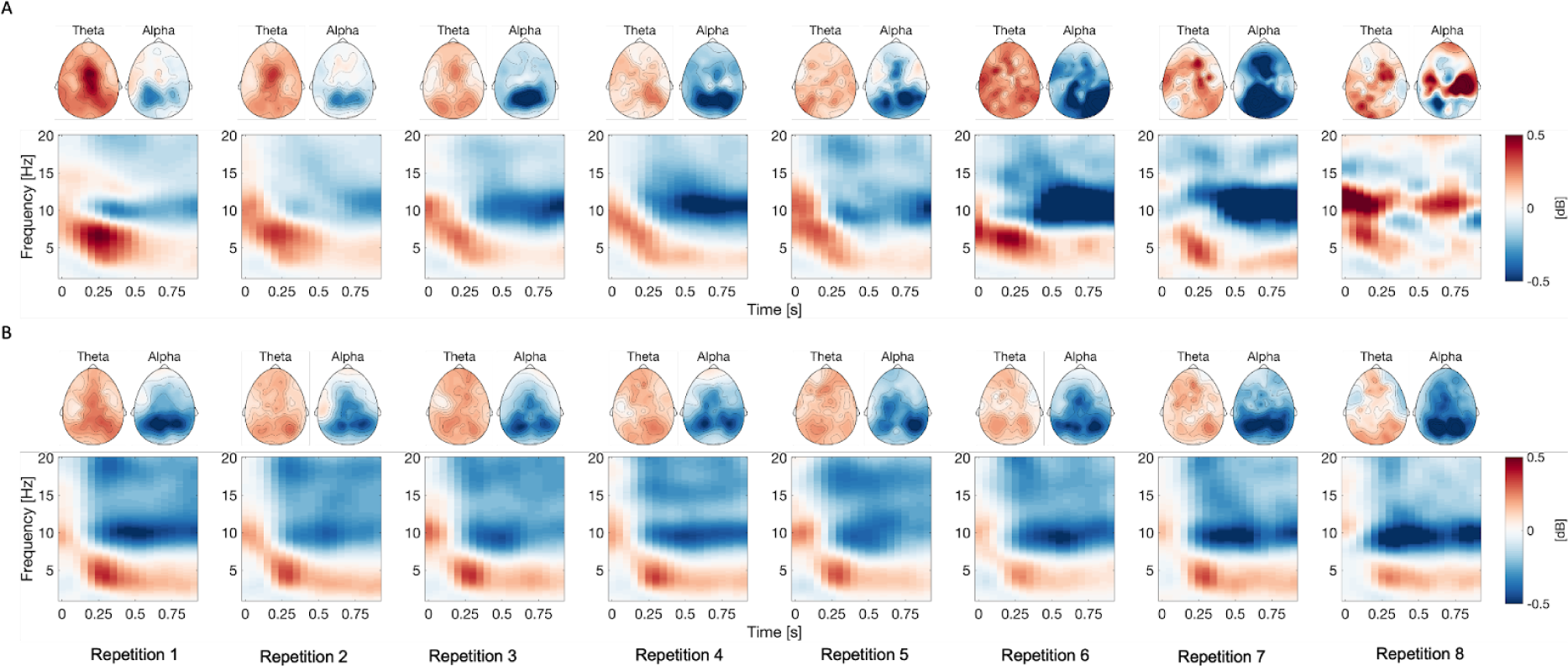
Scalp topographies and time-frequency representations (TFR) of the event-related synchronization (ERS) and desynchronization (ERD) across sequence repetitions over sequence repetitions in young (A) and older (B) subjects. The TFR spectrogram was averaged across all electrodes. Mid-frontal theta and parietal alpha topographies were averaged according to individual alpha frequency (IAF). Blue colors signal event-related desynchronization, red colors signal event-related synchronization. It is important to note that the TFRs of young subjects during repetitions 6, 7, and 8 appear noisy due to the low number of trials.

#### Mid-frontal theta ERS

First, we established the best-fit model for mid-frontal theta ERS. The model included fixed effects of repetition number, age group, and their interaction, and a random effect of subject (Table 2).

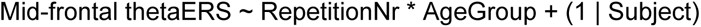

**Table 2.** Effects of repetition number and age group on mid-frontal theta ERS.

| **Variable** | **$\beta$** | **SE** | **CI** | **t-value** | **p-value** |
| --- | --- | --- | --- | --- | --- |
| Intercept | 0.53 | 0.04 | 0.44 – 0.62 | 11.83 | 2.96e-31*** |
| RepetitionNr | -0.09 | 0.01 | -0.12 – -0.07 | -6.55 | 6.48e-11*** |
| AgeGroup [Old] | -0.26 | 0.06 | -0.38 – -0.15 | -4.45 | 9.18e-6*** |
| RepetitionNr*AgeGroup [Old] | 0.08 | 0.02 | 0.04 – 0.11 | 4.61 | 4.19e-6*** |
| **Variance components** | **SD** | **Goodness of fit** |  |  |  |
| Subject | 0.18 | Log likelihood |  | -6947.71 |  |
| Residual | 0.77 |  |  |  |  |

*Note.* Intercept represents the first repetition of young participants. RepetitionNr = Repetition number. $\beta$ = unstandardized regression coefficient. SE = Standard error. CI = Confidence interval. SD = Standard deviation.
\* $p < 0.05$ . \*\* $p < 0.01$ . \*\*\* $p < 0.001$ .

The model revealed a significant main effect of repetition number (β = -0.09; p = 6.48e-11; CI = [-0.12, -0.07]), indicating a decrease of mid-frontal theta ERS with each sequence repetition. Furthermore, there was a significant main effect of age group (β = -0.26; p = 9.18e-6; CI = [-0.38, -0.15]), that is, the mid-frontal theta ERS was decreased in older compared to young subjects. There was also a significant interaction of repetition number and age group (β = 0.08; p = 4.19e-6; CI = [0.04, 0.11]), indicating a smaller decrease of mid-frontal theta ERS across sequence repetitions in older subjects. In addition, we observed a substantial variation of mid-frontal theta ERS between subjects (SD = 0.18).

#### Parietal alpha ERD

Next, we established the best-fit model for parietal alpha ERD. The model included fixed effects of repetition number, age group, and their interaction, and a random effect of subject (Table 3).

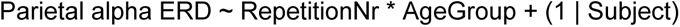

**Table 3.**
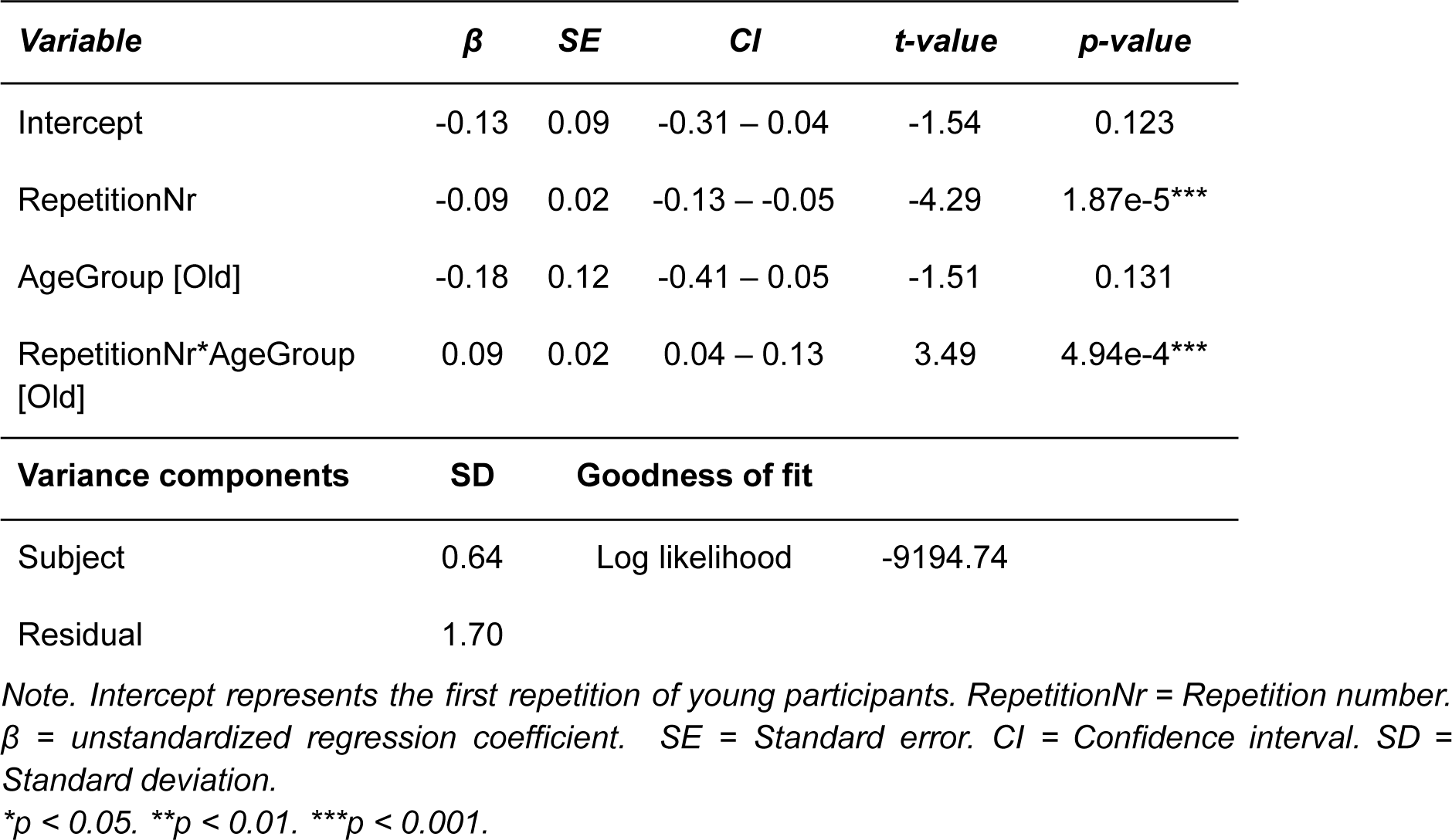
Effects of repetition number and age group on parietal alpha ERD.

Contrary to our hypothesis, the model revealed a significant main effect of repetition number (β = -0.09; p = 1.87e-5; CI = [-0.13, -0.05]), indicating a greater parietal alpha ERD (i.e., increasing desynchronization) with each sequence repetition. The main effect of age group (β = -0.18; p = 0.131; CI = [-0.41, 0.05]) was not statistically significant, providing not enough evidence for age-related differences in overall parietal alpha ERD aggregated across repetitions between young and older subjects. However, there was a significant interaction of repetition number and age group (β = 0.09; p = 4.94e-4; CI = [0.04, 0.13]), reflecting a more gradual increase of parietal alpha ERD across sequence repetitions in older subjects. In addition, we observed a substantial variation of parietal alpha ERD between subjects (SD = 0.64).

In the abovementioned analyses, we employed the specParam algorithm to disentangle periodic and aperiodic components from the EEG signal. As a control analysis and to confirm previous ERS/ERD findings, we also conducted the analysis using conventionally computed power measurements, without the removal of aperiodic component. In summary, our findings reaffirmed that the conventionally computed mid-frontal theta ERS demonstrates a decrease (i.e., reduced synchronization), while parietal alpha ERD exhibits an increase (i.e., increasing desynchronization) over the course of learning. For detailed information please refer to Supplementary Material 5. 3.

So far, we demonstrated that the mid-frontal theta ERS decreases and parietal alpha ERD increases over the course of learning. However, the modulations of mid-frontal theta ERS and parietal alpha ERD could be simply an effect of habituation and the power could decrease or increase as a function of time spent on a task. Therefore, to provide stronger evidence for the relationship of mid-frontal theta ERS and parietal alpha ERD with learning, we conducted a sensitivity analysis to test whether the modulations of both signals change as a function of learning state, that is, from stimulus being unknown, to newly learned, known and finally very well known, regardless of the sequence repetition number. In summary, this analysis demonstrated that mid-frontal theta ERS and parietal alpha ERD exhibit distinct trajectories during the learning process. Specifically, parietal alpha ERD progressively increased (i.e., increasing desynchronization) from trials being unknown, to newly learned and finally fully known. Conversely, mid-frontal theta ERS reached its maximum at the point of first accurate recall (i.e., newly learned), from where it started to gradually decline. Notably, both age groups demonstrated similar patterns in these signals. However, the modulations of mid-frontal theta ERS and alpha ERD were more pronounced in the younger group (Supplementary Material 5. 4).

To further substantiate the connection between both neurophysiological measures (i.e., mid-frontal theta ERS and parietal alpha ERD), and the process of learning, we performed a finer temporal analysis than our earlier learning state categorizations. To this end, we aligned the sequence repetitions to the point where the full sequence was correctly recalled for the first time (i.e., considering the number of repetitions before or after this point). The first correctly recalled sequence was labeled as distance 0, with previous repetitions assigned negative distances and subsequent ones positive distances. The results substantiated our analysis of the learning states. Parietal alpha ERD exhibited a gradual increase (i.e., increasing desynchronization), while mid-frontal theta ERS showed a distinct pattern: initiating with elevated levels, mid-frontal theta ERS exhibited a brief decline, followed by an increase up to the point of the first fully accurate recall (i.e., newly learned), after which it consistently decreased. For detailed insights, please refer to the supplementary material (Supplementary Material 5. 5).

### 3. 3. Mediating effects of pre-stimulus activity on mid-frontal theta ERS and parietal alpha ERD

To assess whether pre-stimulus activity influences the strength of ERS/ERD, we extracted the pre-stimulus activity and used a mediation analysis ^59^ to investigate whether the effect of learning progress (i.e., RepetitionNr) on mid-frontal theta ERS and parietal alpha ERD can be explained by a mediating effect of pre-stimulus power. The premise for a consecutive mediation analysis is a main effect of repetition number in both Model M0 (i.e., effect of repetition number on ERS/ERD) or Model M (i.e., effect of repetition number on pre-stimulus activity). The progression of pre-stimulus and event-related activity is visualized in the figure 6. Both, pre-stimulus theta and alpha power increased over the course of learning. However, the increase was more pronounced in the younger group.

**Figure 6:**
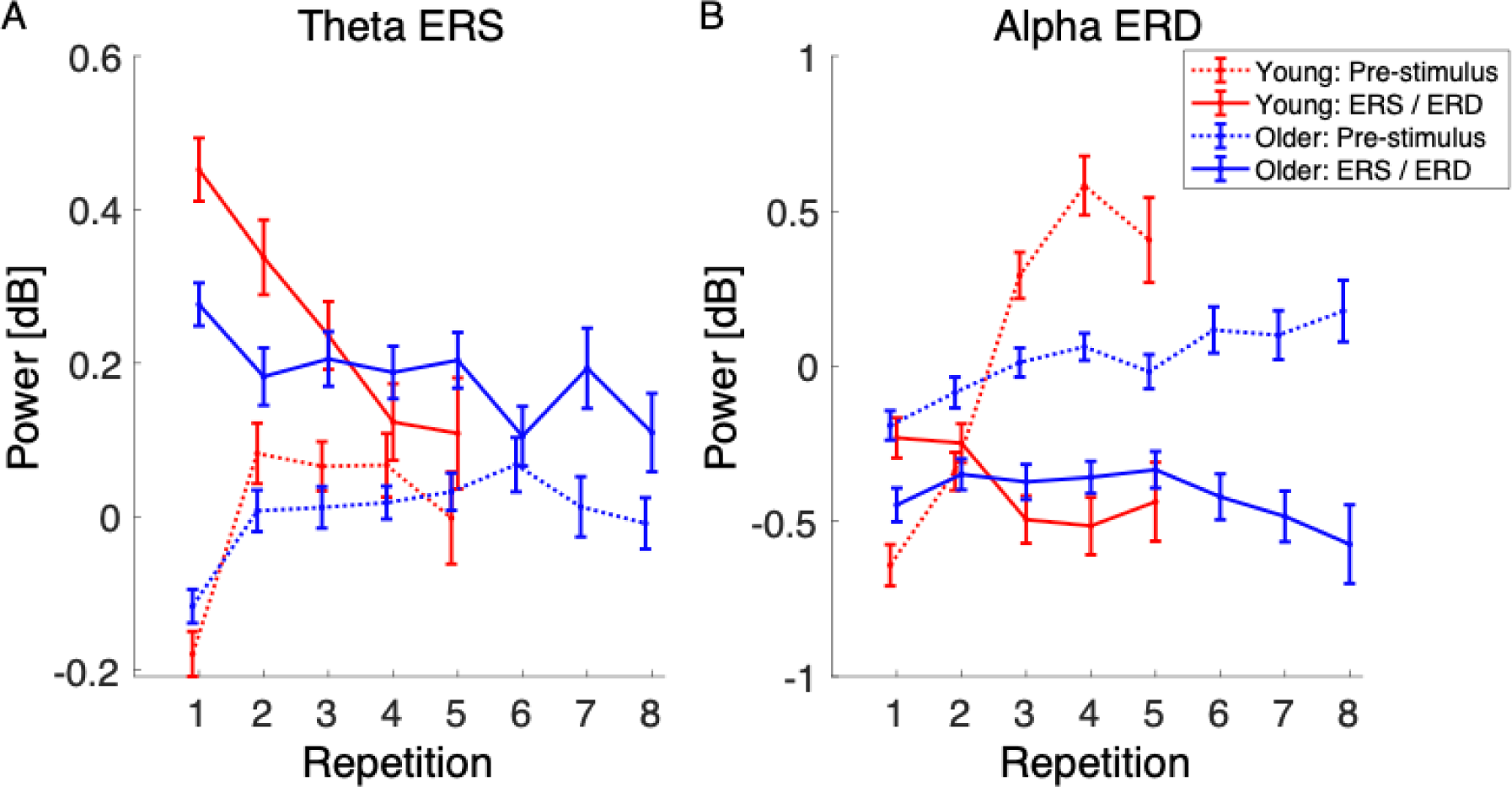
Pre-stimulus and event-related activity over the course of learning in young and older subjects. Please note that some of the subjects memorized the entire sequence within just three repetitions, thus completing the task early. This led to a decrease in the number of subjects, and consequently the data, contributing to repetitions four and beyond. In young, only the first five repetitions were plotted due to the low number of remaining data from the sixth repetition onward. Error bars represent the standard error of the mean.

#### Mid-frontal theta

First, we confirmed a main effect of repetition number on mid-frontal theta ERS (i.e., Model M0) in young (β = -0.19; p = 2e-16; CI = [-0.27, -0.13]) and older (β = -0.04; p = 0.002; CI = [-0.06, -0.01]) subjects, indicating decreasing mid-frontal theta ERS with increasing sequence repetition. Model M revealed that the pre-stimulus mid-frontal theta power increased with sequence repetition in both young (β = 0.14; p = 1.49e-4; CI = [0.07, 0.22]) and older (β = 0.04; p = 8.77e-4; CI = [0.02, 0.07]) subjects. The mediation analysis yielded a significant average causal mediation effect (ACME) of pre-stimulus mid-frontal theta power on mid-frontal theta ERS in young (β = -0.08; p = 2e-16; CI = [-0.12, -0.04]) and older (β = -0.03; p = 2e-16; CI = [-0.04, -0.01]) subjects. An average direct effect (ADE; the effect of repetition number on mid-frontal theta ERS after considering the pre-stimulus mid-frontal theta power) remained significant in young (β = -0.12; p = 2e-16; CI = [-0.18, -0.06]), but not older (β = -0.01; p = 0.202; CI = [-0.03, 0.01]) subjects. Summarized, the total effect of repetition number on mid-frontal theta ERS was partially mediated by pre-stimulus mid-frontal theta power in both age groups (Figure 7).

**Figure 7:**
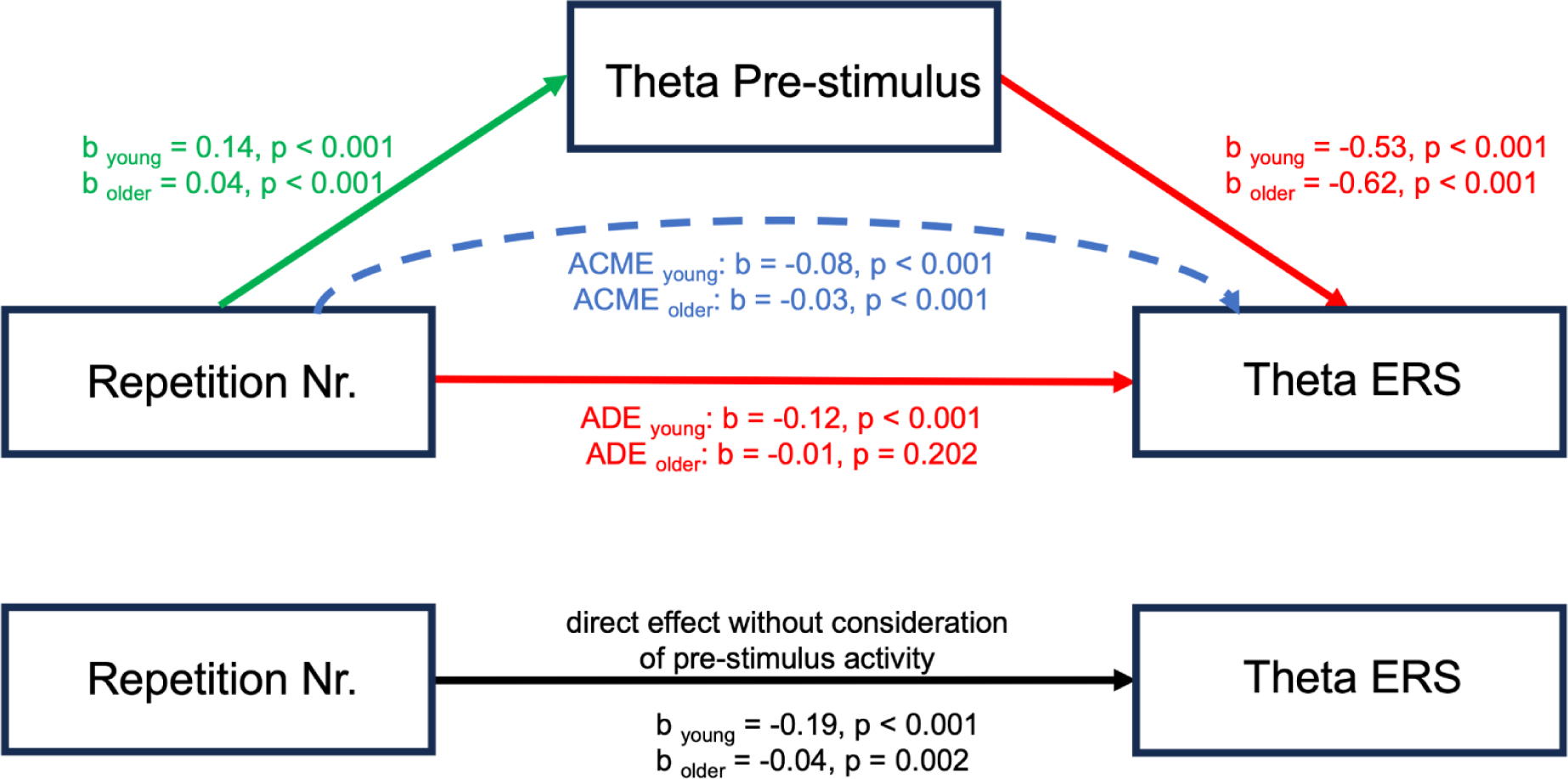
Mediation analysis investigating the effects of pre-stimulus mid-frontal theta power on mid-frontal theta ERS. The results show that a part of the total effect of repetition number on mid-frontal theta ERS is mediated by pre-stimulus mid-frontal theta power in both, young and older subjects. The beta coefficients were standardized to enable direct comparisons.

**Figure 8:**
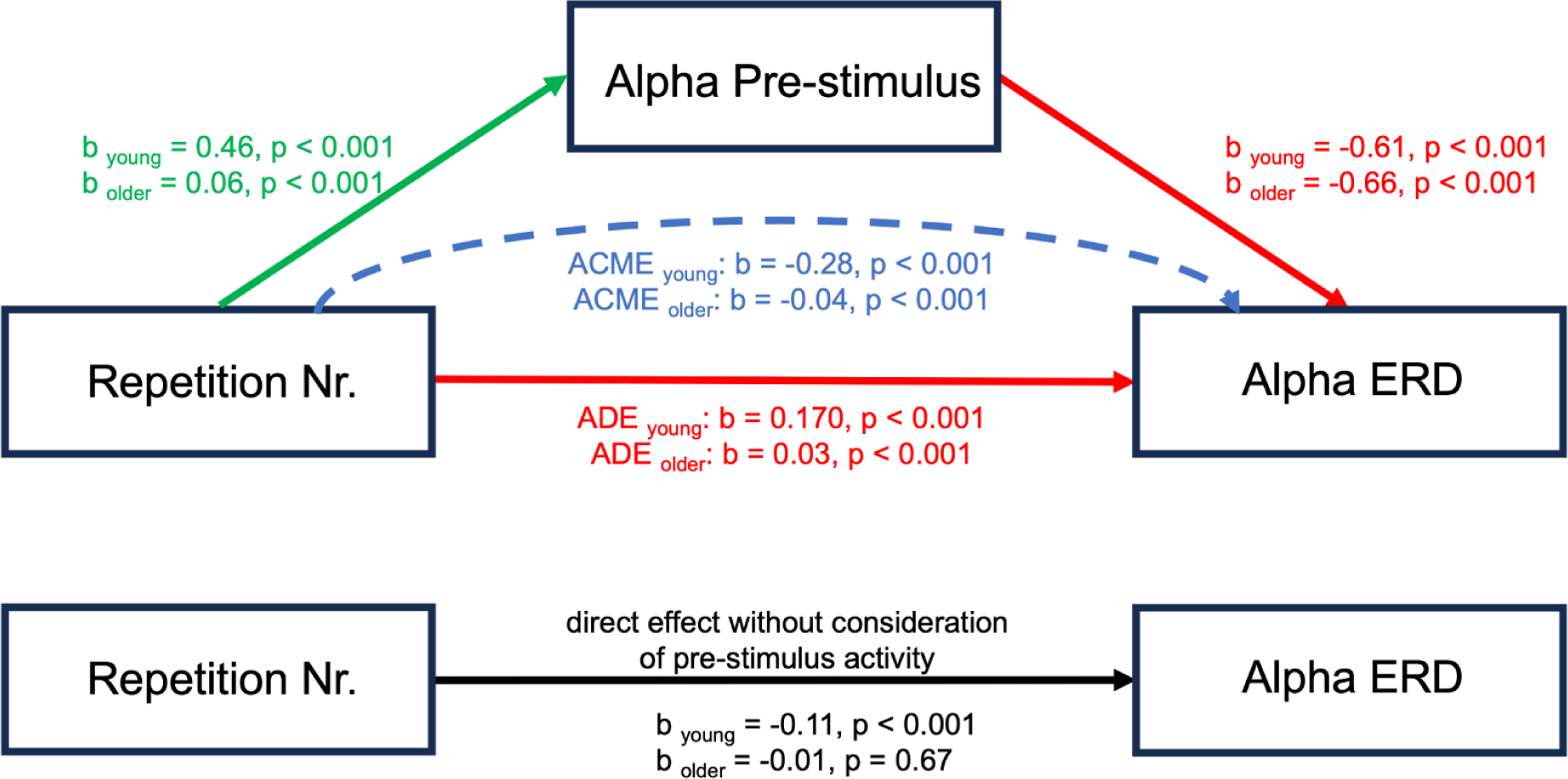
Mediation analysis investigating the effects of pre-stimulus parietal alpha power on parietal alpha ERD. The results show the direct effect (ADE) of repetition number on alpha ERD is positive, whereas the indirect effect (ACME) is negative in both age groups. This suggests that the negative total effect of repetition number on the parietal alpha ERD is entirely due to the influence of the mediator, pre-stimulus parietal alpha. The beta coefficients were standardized to enable direct comparisons.

#### Parietal alpha

Model M0 confirmed the main effect of repetition number on parietal alpha ERD in young subjects (β = -0.11; p = 0.002; CI = [-0.174, -0.04]), indicating greater parietal alpha ERD with increasing sequence repetition. In line with our main analyses, the main effect of repetition number on parietal alpha ERD did not reach significance in older subjects (β = -0.01; p = 0.67; CI = [-0.03, 0.02]), suggesting no significant changes in parietal alpha ERD over sequence repetitions. Model M revealed that the pre-stimulus parietal alpha power increased with sequence repetition in both young (β = 0.46; p = 5.71e-35; CI = [0.39, 0.53]) and older (β = 0.06; p = 1.78e-6; CI = [0.04, 0.08]) subjects. The mediation analysis yielded a significant average causal mediation effect (ACME) of pre-stimulus parietal alpha power on parietal alpha ERD in young (β = -0.28; p = 2e-16; CI = [-0.33, -0.24]) and older (β = -0.04; p = 2e-16 ; CI = [-0.06, -0.02]) subjects. An average direct effect (ADE; the effect of repetition number on parietal alpha ERD after considering the pre-stimulus parietal alpha power) was significant in young (β = 0.17; p = 2e-16; CI = [0.12, 0.22]) and older (β = 0.03; p = 2e-16; CI = [0.02, 0.05]) subjects.

## 4. Discussion

This study investigated the neural processes underlying successful or impeded learning in healthy aging, focusing on event-related mid-frontal theta synchronization (ERS), parietal alpha desynchronization (ERD), as well as on how pre-stimulus activity impacts these neurophysiological indices. Consistent with previous research, we found that younger participants learned faster compared to older participants, highlighting age-related declines in processing speed and working memory (WM) efficiency. Utilizing a visual sequence learning paradigm, our findings revealed the critical mediating role of pre-stimulus activity on mid-frontal theta ERS and parietal alpha ERD. Mid-frontal theta ERS decreased over the course of learning, as the knowledge strengthened, reflecting the decreasing need for mid-frontal theta-mediated memory binding into coherent memory traces. However, causal mediation analysis showed that a significant part of the total mid-frontal theta ERS effect was mediated by the pre-stimulus mid-frontal theta activity. Crucially, the expected decrease in parietal alpha ERD (i.e., reduced desynchronization) throughout the learning process reflecting the general mechanism of efficient information processing, became apparent only after considering pre-stimulus alpha activity. Moreover, significant effects of learning on parietal alpha ERD in older age groups were observed only after accounting for pre-stimulus alpha activity, which would otherwise mask these effects. In the following, we will discuss the implications of these findings for our understanding of age-related changes in neural processing during learning.

### Memory performance decline associated with aging

Consistent with prior research, our behavioral results demonstrated that younger participants learned faster and more efficiently than their older counterparts, as evidenced by higher accuracy, fewer repetitions needed to finish the task, and quicker reaction times (despite the absence of instructions to prioritize speed, see Supplementary Material) ^33,62^. Decline of memory performance in older age is often attributed to biological changes associated with aging, such as decreased neurotransmitter levels and diminished myelination, which can adversely affect neural communication ^63,64^. Van der Linden, Brédart, and Beerten (1994) suggest that attentional resources linked to the frontal cortex undergo a marked decline with aging whereas the storage capacity remains relatively unaffected ^65^. Previously, we demonstrated that the slower and more variable timing of the stimulus identification process, reflected in the event-related potential P300 latency, suggests that despite the older participants engaging the memory formation process to a comparable extent as young participants, there is less time for it to translate the stimulus location information into a solidified memory trace ^37^. This aligns with the cognitive slowing theory, suggesting that slower processing in older participants leads to reduced working memory capacity. This occurs because relevant processes cannot be executed efficiently. Consequently, information is lost at a faster rate during processing in older adults.

### Neurophysiological underpinnings of memory formation in healthy aging

The main goal of this study was to investigate the neural underpinnings contributing to successful or impeded learning in healthy aging. First, we investigated whether observed memory performance deficits were linked either to item binding deficits during encoding, as reflected by attenuated mid-frontal theta event-related synchronization (mid-frontal theta ERS), or to reduction in the depth of information processing, as indexed by parietal alpha event-related desynchronisation (parietal alpha ERD) ^66^.

Our findings revealed that in both age groups, mid-frontal theta ERS decreased as the sequence became more familiar and memory strengthened, indicating a reduction in cognitive load over the course of learning. This is expected because less cognitive effort is needed to process or recall familiar information compared to new information ^6^. The reduction in mid-frontal theta ERS could indicate that less effortful memory encoding was taking place, as less items were required to be bound into coherent memory traces. However, mid-frontal theta ERS was lower and the decrease more gradual in the older group, reflecting more evenly-spread learning progress over time. While some studies report attenuated mid-frontal theta ERS in older age, others find no significant age-related differences, indicating that mid-frontal theta ERS might be necessary but not solely sufficient for the binding of items into coherent memory traces ^11,15,29,30^. Our findings align with the hypothesis that age-related changes in mid-frontal theta ERS play a significant role in the observed differences in associative memory binding during the memory formation process among older adults. Additionally, the application of transcranial direct current stimulation (tDCS) to reduce mid-frontal theta ERS led to slower reaction times compared to sham conditions ^67^. It suggests that mid-frontal theta ERS might directly affect memory encoding efficiency, providing a potential explanation for the age group differences in performance observed in our study.

Contrary to our initial hypothesis, our findings revealed an increase in parietal alpha ERD throughout the learning process in young participants, suggesting a more complex relationship between alpha oscillations and information processing depth than previously anticipated. Additionally, parietal alpha ERD did not change among older participants over the course of learning, indicating limited ability to adapt their information processing strategies and pointing to an overall diminished information processing efficiency.

However, these findings on ERS/ERD require further detailed examination as the process of memory formation depends on both stimulus driven processes and endogenous brain states at the time of stimulus presentation ^68^. To explore the impact of baseline correction on event-related synchronization and desynchronization, we performed a simulation study. We demonstrated that by solely examining the ERS/ERD (as seen in Figure 2 B), it is not possible to definitively determine whether the observed changes in ERS/ERD, throughout the learning process, occurred pre- or post-stimulus. However, when we apply a baseline within the subject and allow the pre-stimulus activity to vary within the subject across sequence repetitions (Figure 2 C), we gain the ability to discern whether the power change occurred pre- or post-stimulus. In essence, this simulation study underscores the critical role of baseline correction in dissecting the temporal dynamics of ERS/ERD and understanding whether changes in brain activity are driven by pre-stimulus or post-stimulus processes. Our study offers novel insight by elucidating the mediating role of pre-stimulus activity in the dynamic relationship between learning progression and mid-frontal theta ERS alongside parietal alpha ERD. For this, we conducted a causal mediation analysis to dissect the contributions of pre-stimulus and post-stimulus activities to the observed relationship between ERS/ERD and behavioral outcomes.

For mid-frontal theta ERS, the mediation analysis yielded a significant average causal mediation effect (ACME) of pre-stimulus mid-frontal theta power on mid-frontal theta ERS in young and older subjects, indicating that a significant part of the total effect of repetition number on mid-frontal theta ERS goes through the pre-stimulus mid-frontal theta power. As the sequence knowledge strengthens, pre-stimulus mid-frontal theta power increases, which in turns lead to decreased mid-frontal theta ERS. The average direct effect (ADE) of learning on mid-frontal theta ERS, which is the effect after consideration of the mediator, pre-stimulus mid-frontal theta, remained significant in young, but not in older subjects. In young subjects, the significant ADE suggests that the modulation of mid-frontal theta ERS is influenced by factors beyond the increase in pre-stimulus mid-frontal theta power. This implies that other processes, such as attention and cognitive control, play a crucial role in mid-frontal theta ERS modulation. In contrast, while the total effect of learning on mid-frontal theta ERS was inherently weaker in older individuals, as indicated by standardized beta estimates, this total effect diminishes to non-significant ADE when accounting for pre-stimulus activity. It suggests that older individuals may exhibit less effective engagement in attention and cognitive control mechanisms, which in turn adversely affects their memory formation capabilities.

For parietal alpha ERD, the mediation analysis revealed what is often termed *inconsistent mediation* ^69^. In this case, the direct effect is found to be positive, while the indirect effect is negative. This scenario implies that the negative impact of the exposure (repetition number) on the outcome (parietal alpha ERD) can be entirely attributed to the influence of the mediator (pre-stimulus alpha). While direct pathways, not involving the mediator, are indeed present in an opposite direction, their influence is not potent enough to offset the considerable impact of the indirect pathways. Crucially, an examination of the total effect in isolation revealed an increase in parietal alpha ERD (indicating greater desynchronization) over the course of learning in young participants, which contradicts our initial hypotheses and the findings derived from the SME paradigm. Given that parietal alpha ERD is associated with the depth of elaborative information processing, one would anticipate the highest parietal alpha ERD levels during the initial repetitions, where the volume of information to be processed is greatest. Subsequently, as knowledge consolidation occurs, parietal alpha ERD levels are expected to decline due to the reduction in novel information requiring processing. Critically, upon separating the contributions of parietal alpha ERD and pre-stimulus activity, and considering the mediating role of pre-stimulus alpha, we observe a significantly positive ADE of learning on parietal alpha ERD across both age groups. This indicates a decrease in parietal alpha ERD (less desynchronization) over the course of learning, aligning with our hypothesis and the existing literature. The observed negative total effect can thus be entirely attributed to the increase in pre-stimulus activity. Moreover, older subjects exhibited a non-significant total effect of learning on parietal alpha ERD but a significant positive ADE and a significant negative ACME. It suggests that while direct cognitive engagement with new information still triggers appropriate neural responses in older adults, the overall effectiveness of these responses is mediated by baseline levels of brain activity, possibly reflecting broader changes in neural efficiency with age. These findings highlight the importance of the mediator, pre-stimulus activity, in explaining the connection between repetition number and parietal alpha ERD. It is evident that the mediator plays a vital role in this relationship, contributing to the intricate dynamics of this mediation process.

Both pre-stimulus mid-frontal theta and parietal alpha power have been associated with successful memory formation. Specifically, pre-stimulus mid-frontal theta has been linked to subsequent encoding of item information ^70–73^ and binding of interlinked information, regardless of stimulus modality ^10^. Furthermore, it was linked to the synchronization of medial prefrontal neurons during learning, facilitating the transfer of hippocampal-dependent traces to neocortical sites ^74^. Previous research suggests that pre-stimulus mid-frontal theta resembles a specific preparatory state for memory processing ^10^, possibly reflecting engagement with a motivational context that influences memory retrieval. However, aging impacts pre-stimulus mid-frontal theta power leading to reduced pre-stimulus mid-frontal theta activity ^75^. Our study contributes to the current literature by demonstrating that pre-stimulus mid-frontal theta power increases over the course of learning in both age groups, with a notably sharper increase in younger participants. This suggests age-related alterations in the capacity to enter preparatory neural states critical for efficient memory encoding and retrieval. On the other hand, higher levels of pre-stimulus parietal alpha power have been linked to neural readiness or idling that can influence subsequent information processing ^31,76–79^. Taken together, pre-stimulus activity can provide insights into anticipatory or preparatory cognitive processes, which might be associated with expectation, attention, or mental readiness. We showed that pre-stimulus alpha power increased over the course of learning in both age groups, however the increase was much stronger in younger subjects. This finding suggests that pre-stimulus alpha activity not only serves as a marker of neural readiness but also differentially impacts the efficacy of learning processes across the lifespan. However, whether pre-stimulus activity reflects general memory-promoting state or it is specifically linked to cognitive processes that are related to memory formation, is still unclear ^10^. Experimental designs that manipulate anticipatory states, such as varying the level of expectation or readiness, would help in disentangling whether pre-stimulus alpha power serves as a general enhancer of cognitive performance or is directly linked to specific memory-related processes.

### Conclusion

Our study provides important insights into the neural underpinnings of age-related differences in learning and memory formation by employing a learning paradigm that captures the incremental nature of learning. We observed a decrease in mid-frontal theta ERS over the course of learning, indicating a reduced reliance on theta-driven mechanisms for integrating memories. This effect, however, was partially mediated by the pre-stimulus mid-frontal theta activity. Importantly, parietal alpha ERD, which reflects efficient information processing, only showed the anticipated decrease over the course of learning after accounting for mediating effects of pre-stimulus parietal alpha activity. Additionally, older subjects showed a non-significant total effect of learning on parietal alpha ERD, but a significant positive direct effect and a significant negative indirect effect. Our study underscores the critical role of pre-stimulus brain states in modulating learning outcomes and elucidating age-related differences. It highlights the need for future research to further investigate the mechanisms underlying the mediating effects of pre-stimulus activity and explore its potential as a target for cognitive enhancement interventions.

## Authorship Contribution

Dawid Strzelczyk: Conceptualization, Investigation, Methodology, Formal analysis, Data curation, Resources, Writing - original draft, Writing - review & editing, Visualization.

Nicolas Langer: Conceptualization, Investigation, Methodology, Formal analysis, Writing - original draft, Writing - review & editing, Visualization, Supervision, Project administration, Resources, Funding acquisition.

## Acknowledgment

This work was supported by the Swiss National Science Foundation.

## Data Availability

All data, preprocessing and analysis scripts used for the analyses are uploaded on OSF.io at https://osf.io/da8k3/. We agree to share our data, any digital study materials, and laboratory logs for all published results in this repository.

