## Supplementary Material for "Pre-stimulus Activity Mediates Event-Related Theta Synchronization and Alpha Desynchronization During Memory Formation in Healthy Aging"

### 1. Supplementary Material

#### 1.1. Reaction times

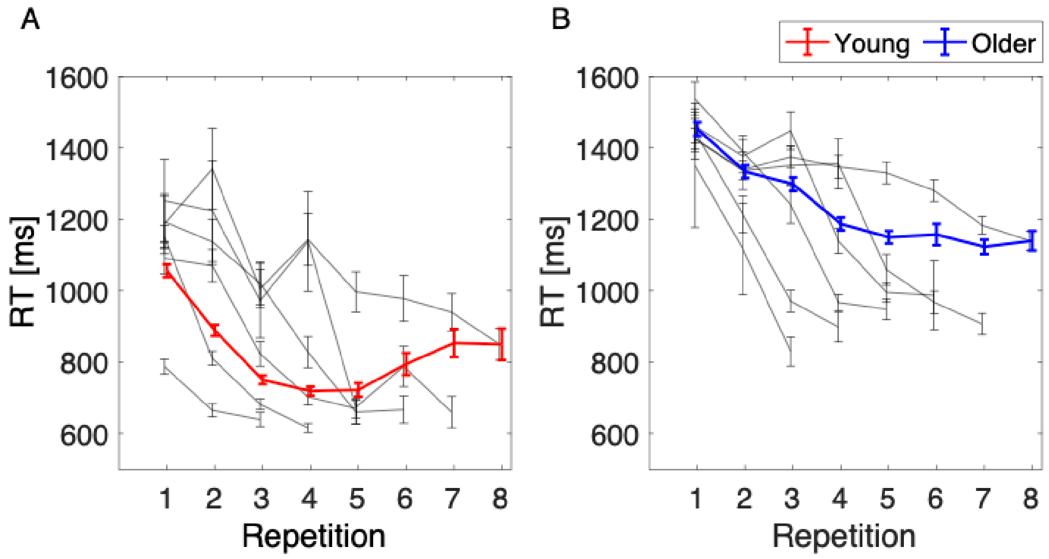


Supplementary Figure 1: Reaction times in young (A) and older (B) participants. Please note that the participants were not explicitly instructed to perform the task as fast as possible. Hence, the reaction times need to be interpreted with caution.

#### 1.2. Grand average time-frequency representation

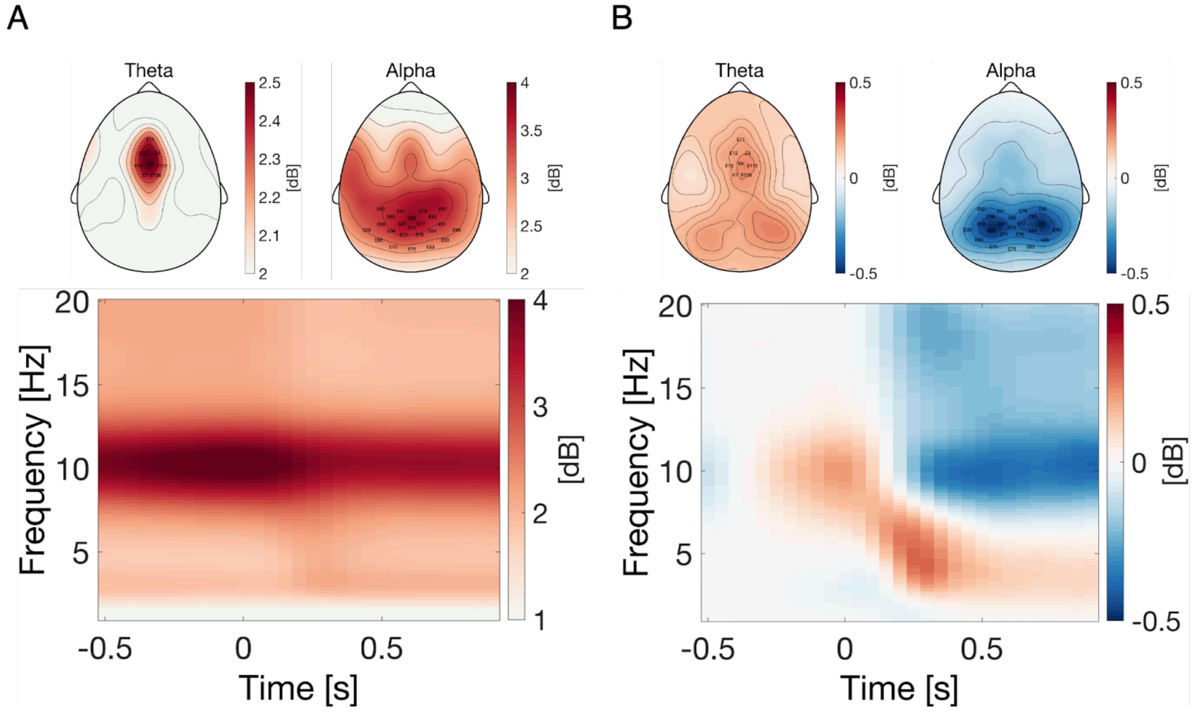


Supplementary Figure 2: Grand average time-frequency representation (TFR) over all age groups, sequence repetitions and channels. (A) Aperiodic adjusted total power. (B) Change in aperiodic-adjusted power relative to pre-stimulus baseline (i.e., -500 to -250 ms). Scalp topographies were averaged over time windows spanning 100 to 500 ms for mid-frontal theta, and 250 to 900 ms for parietal alpha. Highlighted electrodes were used for further statistical analysis.

#### 1.3. Conventionally analyzed ERS/ERD over sequence repetition

In the main manuscript, we employed the specParam algorithm to disentangle periodic and aperiodic components from the EEG signal. As a control analysis and to confirm previous ERS/ERD findings, we also conducted the analysis using conventionally computed power, without the removal of aperiodic components. We followed the same procedure and data analysis steps as in section XX up to aperiodic slope removal. We started the analysis by computing the same model on theta ERS that we previously applied after the removal of aperiodic component.

ThetaERS ~ RepetitionNr * AgeGroup + (1 | ID) + (1|SequenceNr)

The model revealed a significant main effect of repetition number ($\beta = -0.20$; $p = 1.96e\text{-}16$; CI = [-0.25, -0.15]), indicating a decrease of mid-frontal theta ERS with each sequence repetition. Furthermore, there was a significant main effect of age group ($\beta = -0.27$; $p = 0.009$; CI = [-0.48, -0.07]), that is, the mid-frontal theta ERS was decreased in older compared to young subjects. There was also a significant interaction of repetition number and age group ($\beta = 0.16$; $p = 1.36e\text{-}8$; CI = [0.10, 0.21]), indicating a more gradual decrease of mid-frontal theta ERS across sequence repetitions in older subjects (Supplementary Table 1).

Supplementary Table 1. Effects of repetition number and age group on mid-frontal theta ERS

| ***Variable*** | ***β*** | ***SE*** | ***CI*** | ***t-value*** | ***p-value*** |
|---|---|---|---|---|---|
| Intercept | 1.36 | 0.08 | 1.20 – 1.52 | 16.28 | 2.26e-41*** |
| RepetitionNr | -0.20 | 0.02 | -0.25 – -0.15 | -8.25 | 1.96e-16*** |
| AgeGroup [Old] | -0.27 | 0.10 | -0.48 – -0.07 | -2.62 | 0.009** |
| RepetitionNr*AgeGroup [Old] | 0.16 | 0.03 | 0.10 – 0.21 | 5.69 | 1.36e-8*** |
| **Variance components** | **SD** | **Goodness of fit** | | | |
| Subject | 0.41 | Log likelihood | -10561.5 | | |
| SequenceNr | 0.05 | | | | |

| | |
|---|---|
| Residual | 1.49 |

*Note. Intercept represents the first repetition of young participants. RepetitionNr = Repetition number. β = unstandardized regression coefficient. SE = Standard error. CI = Confidence interval. SD = Standard deviation.*
**p < 0.05. **p < 0.01. ***p < 0.001.*

Next, we proceeded to conduct a similar analysis for parietal alpha ERD. Again, we applied the same statistical model as described in the main manuscript.

$$AlphaERD \sim RepetitionNr * AgeGroup + (1 \mid ID)$$

This model revealed a significant main effect of repetition number (β = -0.11; p = 2.99e-11; CI = [-0.14, -0.08]), indicating a greater parietal alpha ERD with each sequence repetition. The main effect of age group (β = -0.40; p = 5.82e-7; CI = [-0.56, -0.25]) was statistically significant, providing evidence for age-related differences in overall parietal alpha ERD aggregated across repetitions between young and older subjects. Furthermore, there was a significant interaction of repetition number and age group (β = 0.10; p = 1.07e-7; CI = [0.06, 0.14]), reflecting a more gradual increase of parietal alpha ERD across sequence repetitions in older subjects (Supplementary Table 2). Summarized, the conventionally computed power revealed a significant main effect of the age group. In contrast, when we analyzed the power with the aperiodic component removed, this age-related effect was not observed. This absence indicates that the influence of age on EEG measurements might predominantly reside within the aperiodic components of the signal, rather than the periodic components.

Supplementary Table 2. Effects of repetition number and age group on parietal alpha ERD

| ***Variable*** | ***β*** | ***SE*** | ***CI*** | ***t-value*** | ***p-value*** |
|---|---|---|---|---|---|
| Intercept | 0.24 | 0.06 | 0.12 – 0.36 | 4.02 | 6.32e-5*** |
| RepetitionNr | -0.11 | 0.02 | -0.14 – -0.08 | -6.66 | 2.99e-11*** |
| AgeGroup [Old] | -0.40 | 0.08 | -0.56 – -0.25 | -5.04 | 5.82e-7*** |
| RepetitionNr*AgeGroup [Old] | 0.10 | 0.02 | 0.06 – 0.14 | 5.32 | 1.07e-7*** |
| **Variance components** | **SD** | **Goodness of fit** | | | |
| Subject | 0.38 | Log likelihood | | -8414.25 | |
| Residual | 1.02 | | | | |

*Note. Intercept represents the first repetition of young participants. RepetitionNr = Repetition number. β = unstandardized regression coefficient. SE = Standard error. CI = Confidence interval. SD = Standard deviation.*

**p* < 0.05. ***p* < 0.01. ****p* < 0.001.*

#### 1.4. ERS/ERD over learning categories

So far, we demonstrated that the mid-frontal theta ERS decrease and parietal alpha ERD increase with sequence repetitions. However, the modulations of mid-frontal theta and parietal alpha could be simply an effect of habituation and the power could decrease or increase as a function of time spent on a task. Therefore, to provide stronger evidence for the relationship of mid-frontal theta ERS and parietal alpha ERD with learning, we tested whether the modulations of both components change as a function of learning state, that is, from stimulus being unknown, to newly learned, known and finally very well known, regardless of the sequence repetition number.

First, each individual stimulus presentation was classified as one of five categories based on the responses of the participants after each sequence repetition. A stimulus was assigned to the *unknown* category (UN) when location was recalled incorrectly in the current and all previous repetitions, to the *newly learned* category (NL) when stimulus position was recalled correctly for the first time in the current repetition, to the *known* category (K) when stimulus position was correctly recalled at least twice in a row, to the *very well known* category (K+) when stimulus position was correctly recalled at least three times in a row, and to the *forgotten* category (F) when stimulus position was once recalled correctly but incorrectly in the following sequence repetition. The *forgotten* category was removed from further analysis due to the low number of trials. Next, we identified the best-fit model for the mid-frontal theta ERS and parietal alpha ERD. Because we were merely interested in power differences across learning categories and age groups, we additionally computed contrasts for those variables using the *emmeans* package in R Studio [80]. Supplementary Figure 3 shows the time-frequency decomposition of young and older subjects averaged over all 105 electrodes as a function of the learning category.

#### Mid-frontal theta

$$\text{ThetaERS} \sim \text{Category} * \text{AgeGroup} + (1 \mid \text{ID})$$

The model revealed a significant main effect of category K+ ($\beta = -0.19$; $p = 0.010$ ; CI = [-0.33, -0.05]), indicating decreased mid-frontal theta ERS for K+ compared to UN (i.e., intercept) trials. The other effects did not reach significance (Supplementary Table 3). Next, we proceeded with computing contrasts between learning categories and age groups. The results of contrast comparisons are summarized in Supplementary Table 4.

Supplementary Table 3. Effects of learning category and age group on mid-frontal theta ERS

| ***Variable*** | ***β*** | ***SE*** | ***CI*** | ***t-value*** | ***p-value*** |
|---|---|---|---|---|---|
| Intercept | 0.34 | 0.07 | 0.21 – 0.46 | 5.14 | 2.81e-7*** |
| NL | 0.12 | 0.07 | -0.02 – 0.26 | 1.63 | 0.103 |
| K | -0.07 | 0.07 | -0.21 – 0.07 | -0.95 | 0.342 |
| K+ | -0.19 | 0.07 | -0.33 – -0.05 | -2.60 | 0.010* |
| AgeGroup [Old] | -0.12 | 0.08 | -0.27 – 0.04 | -1.49 | 0.136 |
| NL*AgeGroup [Old] | -0.11 | 0.09 | -0.29 – 0.07 | -1.16 | 0.247 |
| K*AgeGroup [Old] | 0.06 | 0.09 | -0.13 – 0.24 | 0.60 | 0.550 |
| K+*AgeGroup [Old] | 0.12 | 0.09 | -0.05 – 0.31 | 1.45 | 0.147 |
| **Variance components** | **SD** | **Goodness of fit** | | | |
| Subject | 0.17 | Log likelihood | | -4636.92 | |
| Residual | 0.82 | | | | |

*Note. Intercept represents the first repetition of young participants. NL = newly learned, K = known, K+ = better known. β = unstandardized regression coefficient. SE = Standard error. CI = Confidence interval. SD = Standard deviation.*
**p < 0.05. **p < 0.01. ***p < 0.001.*

In young participants, the contrast comparison revealed a slightly larger mid-frontal theta ERS for UN trials relative to K (β = 0.07; p = 0.999; CI = [-0.07, 0.21]) and K+ (β = ; p = ; CI = []) trials, however the differences did not reach statistical significance. Mid-frontal Theta ERS for NL trials was slightly larger than UN trials (β = -0.11; p = 0.927; CI = [-0.25, 0.02]), but this was not statistically significant. NL trials showed significantly larger mid-frontal theta ERS compared to K (β = 0.18; p = 0.001; CI = [0.09, 0.28]) and K+ (β = 0.30; p = 1e-10; CI = [0.21, 0.40]) trials. Finally, K trials exhibited greater mid-frontal theta ERS than K+ trials, but the difference was not statistically significant.

In older subjects, we observed trends similar to those seen in the younger cohort. The mid-frontal theta ERS in UN trials appeared slightly larger when compared to both K (β = 0.01; p = 0.999; CI = [-0.10, 0.13]) and K+ (β = 0.05; p = 0.999; CI = [-0.05, 0.16]) trials. Additionally, NL trials seemed to exhibit a marginally higher mid-frontal theta ERS compared to UN (β = -0.01; p = 0.999; CI = [-0.12, 0.10]) trials. Finally, K trials exhibited greater mid-frontal theta ERS than K+ (β = 0.04; p = 0.999; CI = [-0.04, 0.17]) trials. These trends mirrored the patterns we noted in young participants. However, none of these comparisons in the older group reached statistical significance.

When comparing both age groups, young showed a higher mid-frontal theta ERS in UN trials compared to their older counterparts ($\beta = 0.12$; $p = 0.999$; CI = [-0.03, 0.27]). However, this difference was not statistically significant. Notably, the most pronounced difference was observed in NL trials ($\beta = 0.29$; $p = 1e-6$; CI = [0.18, 0.40]), where young subjects exhibited significantly larger mid-frontal theta ERS than older subjects. Conversely, while young participants also displayed slightly larger mid-frontal theta ERS in K trials ($\beta = 0.06$; $p = 0.999$; CI = [-0.05, 0.17]), this difference did not reach statistical significance. Finally, in K+ trials, young subjects had marginally lower mid-frontal theta ERS compared to older subjects, but again, this was not statistically significant ($\beta = -0.01$; $p = 0.999$; CI = [-0.12, 0.09]).

Supplementary Table 4. Mid-frontal theta ERS: contrast comparisons between age groups and learning categories.

| **Contrast** | ***β*** | ***SE*** | ***CI*** | ***z-ratio*** | ***p-value*** |
|---|---|---|---|---|---|
| UN Young - NL Young | -0.11 | 0.07 | -0.25 – 0.02 | -1.63 | 0.927 |
| UN Young - K Young | 0.07 | 0.07 | -0.07 – 0.21 | 0.95 | 0.999 |
| UN Young - K+ Young | 0.18 | 0.07 | 0.05 – 0.33 | 2.60 | 0.085 |
| NL Young - K Young | 0.18 | 0.05 | 0.09 – 0.28 | 3.85 | 0.001** |
| NL Young - K+ Young | 0.30 | 0.05 | 0.21 – 0.40 | 6.35 | 1e-10*** |
| K Young - K+ Young | 0.11 | 0.05 | 0.02 – 0.21 | 2.45 | 0.129 |
| UN Old - NL Old | -0.01 | 0.05 | -0.12 – 0.10 | 0.86 | 0.999 |
| UN Old - K Old | 0.01 | 0.06 | -0.10 – 0.13 | 0.22 | 0.999 |
| UN Old - K+ Old | 0.05 | 0.06 | -0.05 – 0.16 | 0.98 | 0.999 |
| NL Old - K Old | 0.02 | 0.06 | -0.08 – 0.13 | 0.43 | 0.999 |
| NL Old - K+ Old | 0.06 | 0.05 | -0.04 – 0.17 | 1.23 | 0.999 |
| K Old - K+ Old | 0.04 | 0.05 | -0.04 – 0.17 | 0.22 | 0.999 |
| UN Young - UN Old | 0.12 | 0.08 | -0.04 – 0.27 | 1.49 | 0.999 |
| NL Young - NL Old | 0.29 | 0.05 | 0.18 – 0.40 | 5.35 | 1e-6*** |
| K Young - K Old | 0.06 | 0.06 | -0.05 – 0.17 | 1.12 | 0.999 |
| K+ Young - K+ Old | -0.01 | 0.05 | -0.12 – 0.09 | -0.23 | 0.999 |

*Note. UN = Unknown. NL = Newly learned. K = Known, K+ = Better known. β = unstandardized*

*regression coefficient. SE = Standard error. CI = Confidence interval. P-values adjusted using the Bonferroni method for 16 tests.*
**p < 0.05. **p < 0.01. ***p < 0.001.*

#### Parietal alpha

The model revealed a significant main effect of category K+ (β = -0.44; p = 3e-4; CI = [-0.65, -0.23]), indicating increased parietal alpha ERD for K+ compared to UN (i.e., intercept) trials. Moreover, there was a significant interaction of the K+ trials and age group (β = 0.43; p = 0.001; CI = [0.17, 0.69]), that is older subjects had decreased parietal alpha ERD for K+ trials than young subjects. The other effects did not reach significance (Supplementary Table 5). Next, we proceeded with computing contrasts between learning categories and age groups. The results of contrast comparisons are summarized in Supplementary Table 6.

Supplementary Table 5. Effects of learning category and age group on parietal alpha ERD

| ***Variable*** | ***β*** | ***SE*** | ***CI*** | ***t-value*** | ***p-value*** |
|---|---|---|---|---|---|
| Intercept | -0.10 | 0.11 | -0.31 – 0.12 | -0.86 | 0.392 |
| NL | -0.15 | 0.11 | -0.36 – 0.06 | -1.39 | 0.163 |
| K | -0.18 | 0.11 | -0.39 – 0.03 | -1.66 | 0.097 |
| K+ | -0.44 | 0.11 | -0.65 – -0.23 | -4.15 | 3e-4*** |
| AgeGroup [Old] | -0.19 | 0.14 | -0.47 – 0.08 | -1.37 | 0.170 |
| NL*AgeGroup [Old] | 0.00 | 0.14 | -0.27 – 0.27 | 0.01 | 0.993 |
| K*AgeGroup [Old] | 0.08 | 0.14 | -0.19 – 0.35 | 0.58 | 0.561 |
| K+*AgeGroup [Old] | 0.43 | 0.13 | 0.17 – 0.69 | 3.20 | 0.001** |
| **Variance components** | **SD** | **Goodness of fit** | | | |
| Subject | 0.64 | Log likelihood | | -6196.78 | |
| Residual | 1.20 | | | | |

*Note. Intercept represents the first repetition of young participants. NL = newly learned, K = known, K+ = better known. β = unstandardized regression coefficient. SE = Standard error. CI = Confidence interval. SD = Standard deviation.*
**p < 0.05. **p < 0.01. ***p < 0.001.*

In young participants, the contrast comparison revealed smaller parietal alpha ERD for UN trials relative to NL (β = 0.15; p = 0.999; CI = [-0.06, 0.36]), K (β = 0.18; p = 0.868; CI = [-0.03, 0.39]) and K+ (β = 0.44; p = 2e-4; CI = [0.23, 0.65]) trials, however the difference between UN and K, and UN and NL trials did not reach statistical significance. NL trials exhibited a slightly smaller parietal alpha ERD compared to K trials, though the difference was not statistically significant (β = 0.03; p = 0.999; CI = [-0.11, 0.17]). In contrast, NL trials demonstrated a significantly smaller parietal alpha ERD than K+ (β = 0.29; p = 3e-4; CI = [0.15, 0.43]) trials. Finally, K trials exhibited significantly greater smaller ERD than K+ trials (β = 0.26; p =0.001; CI = [0.13, 0.40]). Summarized, parietal alpha ERD increased as the knowledge strengthened.

Unlike in younger participants where a trend was observed in parietal alpha ERD as knowledge increased, older subjects did not show any significant trends across different learning categories. UN trials showed smaller parietal alpha ERD compared to NL (β = 0.15; p = 0.747; CI = [-0.02, 0.31]), K (β = 0.10; p = 0.999; CI = [-0.07, 0.26]), and K+ (β = 0.01; p = 0.999; CI = [-0.15, 0.17]) trials. However, none of these differences reached statistical significance. NL trials exhibited larger parietal alpha ERD compared to both K (β = -0.05; p = 0.999; CI = [-0.21, 0.11]) and K+ (β = -0.14; p = 0.729; CI = [-0.29, 0.02]) trials, but these differences were not statistically significant. Similarly, K trials had larger parietal alpha ERD compared to K+ (β = -0.09; p = 0.999; CI = [-0.24, 0.06]) trials, yet this difference was also not statistically significant.

Finally, we compared both age groups. The parietal alpha ERD was smaller in young participants compared to older participants in UN (β = 0.19; p = 0.999; CI = [-0.08, 0.47]), NL (β = 0.19; p = 0.828; CI = [-0.03, 0.42]) and K (β = 0.12; p = 0.999; CI = [-0.11, 0.34]) trials. However, none of these effects reached statistical significance. Interestingly, the pattern reversed in K+ trials, where older participants demonstrated smaller, yet not significant parietal alpha ERD compared to young participants (β = -0.23; p = 0.329; CI = [-0.45, -0.01]).

Supplementary Table 6. Parietal alpha ERD: contrast comparisons between age groups and learning categories.

| **Contrast** | ***β*** | ***SE*** | ***CI*** | ***z-ratio*** | ***p-value*** |
|---|---|---|---|---|---|
| UN Young - NL Young | 0.15 | 0.11 | -0.06 – 0.36 | 1.39 | 0.999 |
| UN Young - K Young | 0.18 | 0.11 | -0.03 – 0.39 | 1.66 | 0.868 |
| UN Young - K+ Young | 0.44 | 0.11 | 0.23 – 0.65 | 4.15 | 2e-4*** |
| NL Young - K Young | 0.03 | 0.07 | -0.11 – 0.17 | 0.41 | 0.999 |
| NL Young - K+ Young | 0.29 | 0.07 | 0.15 – 0.43 | 4.18 | 3e-4*** |
| K Young - K+ Young | 0.26 | 0.07 | 0.13 – 0.40 | 3.74 | 0.001** |
| UN Old - NL Old | 0.15 | 0.08 | -0.02 – 0.31 | 1.73 | 0.747 |

| | | | | | |
|---|---|---|---|---|---|
| UN Old - K Old | 0.10 | 0.08 | -0.07 – 0.26 | 1.15 | 0.999 |
| UN Old - K+ Old | 0.01 | 0.08 | -0.15 – 0.17 | 0.15 | 0.999 |
| NL Old - K Old | -0.05 | 0.08 | -0.21 – 0.11 | -0.61 | 0.999 |
| NL Old - K+ Old | -0.14 | 0.08 | -0.29 – 0.02 | -1.75 | 0.729 |
| K Old - K+ Old | -0.09 | 0.08 | -0.24 – 0.06 | -1.11 | 0.999 |
| UN Young - UN Old | 0.19 | 0.14 | -0.08 – 0.47 | 1.37 | 0.999 |
| NL Young - NL Old | 0.19 | 0.11 | -0.03 – 0.42 | 1.69 | 0.828 |
| K Young - K Old | 0.12 | 0.12 | -0.11 – 0.34 | 1.00 | 0.999 |
| K+ Young - K+ Old | -0.23 | 0.11 | -0.45 – -0.01 | -2.09 | 0.329 |

*Note. UN = Unknown. NL = Newly learned. K = Known, K+ = Better known. β = unstandardized regression coefficient. SE = Standard error. CI = Confidence interval. P-values adjusted using the Bonferroni method for 16 tests.*
**p < 0.05. **p < 0.01. ***p < 0.001.*

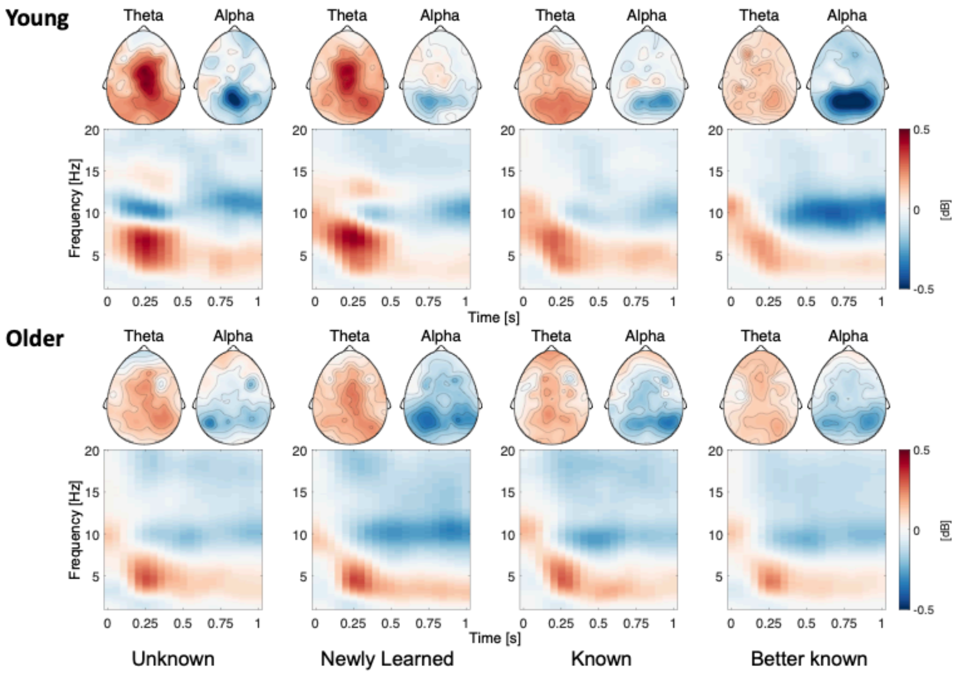


Supplementary Figure 3: Scalp topographies and time-frequency representation (TFR) of the ERS/ERD over learning categories in young (top) and older (bottom) subjects. The TFR spectrogram was averaged across all electrodes. Mid-frontal theta and parietal alpha topographies were averaged according to IAF. Blue colors signal event-related desynchronization (ERD), red colors signal event-related synchronization (ERS).

In summary, mid-frontal theta ERS and parietal alpha ERD exhibit distinct trajectories during the learning process. Parietal alpha ERD displayed a continuous decrease, whereas mid-frontal theta ERS peaked at the point of first accurate recall (i.e., newly learned). Notably, both age groups demonstrated similar patterns in these signals. However, the modulations of mid-frontal theta and parietal alpha were more pronounced in the younger group.

#### 1.5. Finer temporal analysis: distance to first accurate recall

To further substantiate the connection between mid-frontal theta ERS, parietal alpha ERD, and the process of learning, we aligned the sequence repetitions to the point where the entire sequence was correctly recalled for the first time (i.e., considering the number of repetitions before or after this point). The first entire correctly recalled sequence was labeled as distance 0, with previous repetitions assigned negative distances and subsequent ones positive distances. This approach allowed for a more detailed investigation of the learning process over time, offering finer temporal resolution than our earlier learning state categorizations (Supplementary Figure 4).

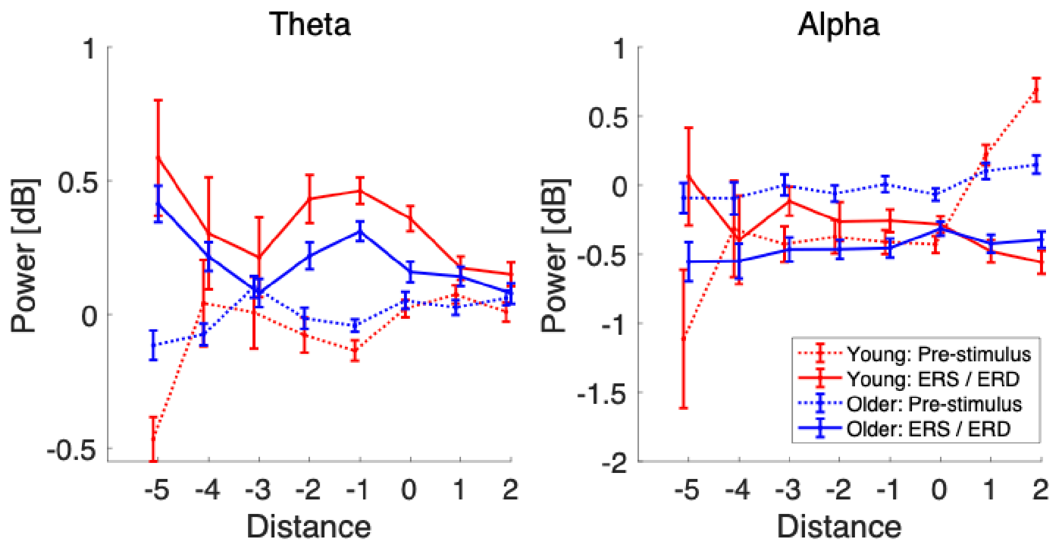


Supplementary Figure 4: Prestimulus power, mid-frontal theta ERS and parietal alpha ERD aligned to the point of first fully accurate recall (i.e., distance of 0). Please note that some of the subjects memorized the entire sequence within just three repetitions (i.e., distances 0, 1, 2), thus completing the task early. This led to a decrease in the number of subjects, and consequently the data, contributing to distances below zero and above two. Error bars represent the standard error of the mean.

Supplementary Table 7. Effects of distance and age group on mid-frontal theta ERS

| ***Variable*** | ***β*** | ***SE*** | ***CI*** | ***t-value*** | ***p-value*** |
|---|---|---|---|---|---|
| Intercept | 0.31 | 0.03 | 0.26 – 0.36 | 11.75 | 3.95e-25*** |
| Distance | -0.06 | 0.01 | -0.08 – -0.04 | -5.35 | 9.22e-8*** |
| AgeGroup [Old] | -0.12 | 0.04 | -0.19 – -0.05 | -3.23 | 0.001** |
| Distance*AgeGroup [Old] | 0.05 | 0.01 | 0.02 – 0.07 | 3.39 | 0.001** |
| **Variance components** | **SD** | **Goodness of fit** | | | |
| Subject | 0.18 | Log likelihood | | -6280.21 | |
| Residual | 0.90 | | | | |

*Note. Intercept represents the distance '-5' of young participants. β = unstandardized regression coefficient. SE = Standard error. CI = Confidence interval. SD = Standard deviation.*

Supplementary Table 8. Effects of distance and age group on parietal alpha ERD

| ***Variable*** | ***β*** | ***SE*** | ***CI*** | ***t-value*** | ***p-value*** |
|---|---|---|---|---|---|
| Intercept | -0.36 | 0.07 | -0.49 – -0.23 | -5.31 | 2.64e-7*** |
| Distance | -0.06 | 0.02 | -0.09 – -0.02 | -3.24 | 0.001** |
| AgeGroup [Old] | -0.00 | 0.10 | -0.19 – 0.19 | -0.02 | 0.985 |
| Distance*AgeGroup [Old] | 0.06 | 0.02 | 0.02 – 0.11 | 2.88 | 0.004** |
| **Variance components** | **SD** | **Goodness of fit** | | | |
| Subject | 0.63 | Log likelihood | | -8357.51 | |
| Residual | 1.37 | | | | |

*Note. Intercept represents the distance '-5' of young participants. β = unstandardized regression coefficient. SE = Standard error. CI = Confidence interval. SD = Standard deviation.*

Supplementary Table 9. Effects of distance and age group pre-stimulus mid-frontal theta ERS

| ***Variable*** | ***β*** | ***SE*** | ***CI*** | ***t-value*** | ***p-value*** |
|---|---|---|---|---|---|
| Intercept | 0.00 | 0.00 | -0.06 – 0.07 | 0.10 | 0.919 |
| Distance | 0.01 | 0.01 | 0.00 – 0.01 | 2.89 | 0.004** |

| Variance components | SD | Goodness of fit | |
|---|---|---|---|
| SequenceNr | 0.08 | Log likelihood | -5033.7 |
| Residual | 0.70 | | |

*Note. Intercept represents the distance '-5' of young participants. SequenceNr = Sequence number. β = unstandardized regression coefficient. SE = Standard error. CI = Confidence interval. SD = Standard deviation.*

Supplementary Table 10. Effects of distance and age group on pre-stimulus parietal alpha ERD

| ***Variable*** | ***β*** | ***SE*** | ***CI*** | ***t-value*** | ***p-value*** |
|---|---|---|---|---|---|
| Intercept | -0.19 | 0.10 | -0.40 – 0.01 | -1.86 | 0.063 |
| Distance | 0.21 | 0.02 | 0.18 – 0.25 | 11.78 | 1.41e-31*** |
| AgeGroup [Old] | 0.17 | 0.05 | 0.07 – 0.26 | 3.48 | 4.97e-4*** |
| ET | 0.02 | 0.01 | 0.01 – 0.04 | 2.56 | 0.010* |
| Distance*AgeGroup [Old] | -0.17 | 0.02 | -0.21 – -0.13 | -7.62 | 2.97e-14*** |
| **Variance components** | **SD** | **Goodness of fit** | | | |
| SequenceNr | 0.23 | Log likelihood | -8558.39 | | |
| Residual | 1.48 | | | | |

*Note. Intercept represents the distance '-5' of young participants. SequenceNr = Sequence number. β = unstandardized regression coefficient. SE = Standard error. CI = Confidence interval. SD = Standard deviation.*

#### 1.6. Moderated mediation

To investigate the age-related variations in how repetition number influences mid-frontal theta ERS and parietal alpha ERD, and to understand the role of pre-stimulus activity in this relationship, we employed a moderated mediation analysis with the age group as a moderator.

#### Mid-frontal theta

Initially, we established a significant main effect of repetition number on mid-frontal theta ERS in young subjects (β = -0.19; p = 2e-16; CI = [-0.25, -0.14]), which demonstrated a decrease in mid-frontal theta ERS with increasing sequence repetition. Subsequently, our Model M indicated that pre-stimulus mid-frontal theta power in young subjects increased with sequence repetition (β = 0.14; p = 3.81e-6; CI = [0.08, 0.20]). However, these effects were found to be moderated by age group, as evidenced by significant interactions between repetition number and the age group. In the case of older subjects, the decrease in mid-frontal theta ERS over repetitions was less pronounced (β = 0.16; p = 4.19e-6; CI = [0.09, 0.23]), and there was a less substantial increase in pre-stimulus mid-frontal theta power (β = -0.10; p = 0.005; CI = [-0.17, -0.03]). The mediation analysis yielded a significant average causal mediation effect (ACME) of pre-stimulus mid-frontal theta power on mid-frontal theta ERS in young (β = -0.08; p = 2e-16; CI = [-0.11, -0.04]) and older (β = -0.03; p = 0.002; CI = [-0.05, -0.01]) subjects. An average direct effect (ADE; the effect of repetition number on mid-frontal theta ERS after considering the pre-stimulus mid-frontal theta power) remained significant in young (β = -0.12; p = 2e-16; CI = [-0.17, -0.07]), but not older (β = 0.01; p = 0.202; CI = [-0.04, 0.01]) subjects (Supplementary Figure 4).

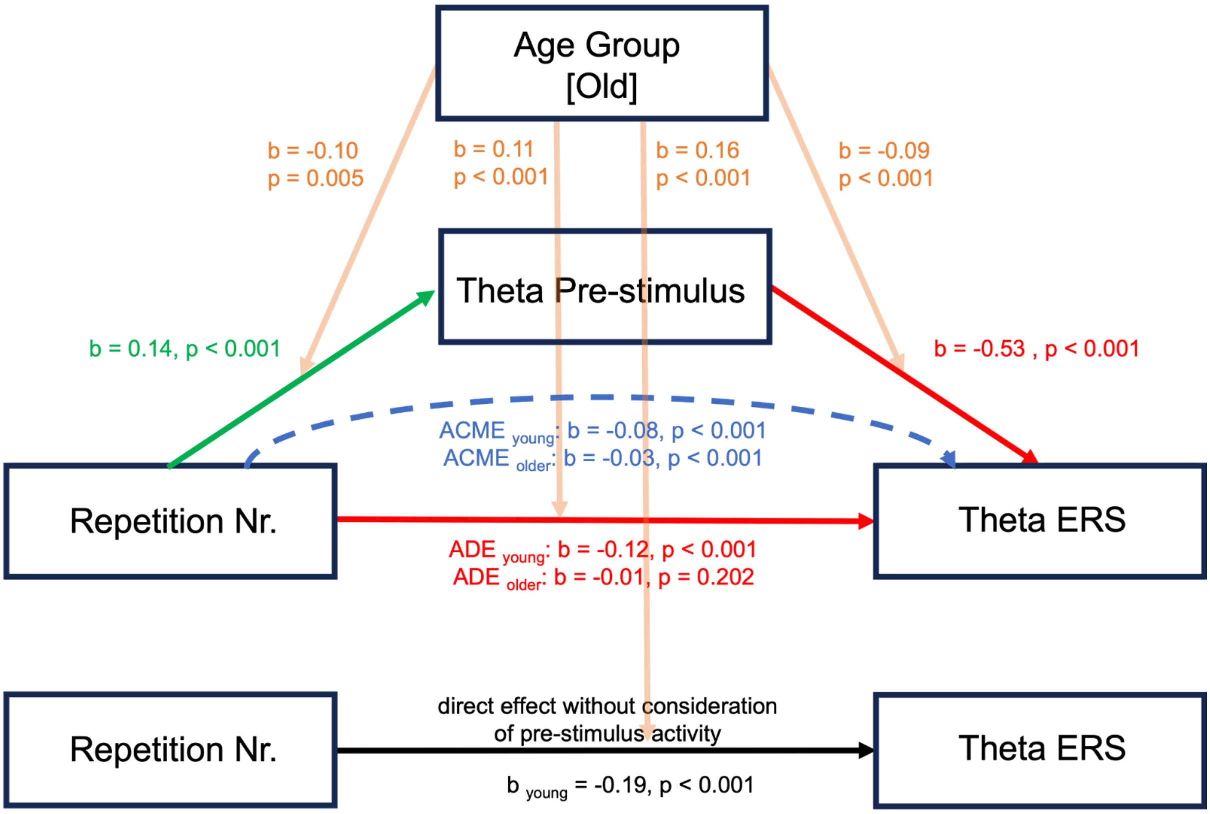


Supplementary Figure 4. Moderated mediation analysis investigating the effects of pre-stimulus mid-frontal theta power on mid-frontal theta ERS. The results show that a part of the total effect of repetition number on mid-frontal theta ERS is mediated by pre-stimulus mid-frontal theta power.

#### Parietal alpha

First, we confirmed a significant main effect of repetition number on parietal alpha ERD in young subjects (β = -0.11; p = 2e-16; CI = [-0.16, -0.06]), which indicated a greater parietal alpha ERD with an increase in sequence repetition. Subsequently, Model M demonstrated that pre-stimulus parietal alpha power in young subjects increased alongside sequence repetition (β = 0.46; p = 4.64e-52; CI = [0.40, 0.52]). However, these effects appeared to be moderated by age group, as shown by significant interactions between repetition number and the age group. For older subjects, the increase in parietal alpha ERD over repetitions was less pronounced (β = 0.11; p = 4.94e-4; CI = [0.05, 0.17]), and there was a more modest increase in pre-stimulus parietal alpha power (β = -0.40; p = 1.51e-31; CI = [-0.47, -0.33]). The mediation analysis yielded a significant average causal mediation effect (ACME) of pre-stimulus parietal alpha power on parietal alpha ERD in both young (β = -0.28; p = 2e-16; CI = [-0.32, -0.25]) and older (β = -0.04; p = 2e-16; CI = [-0.06, -0.02]) subjects. Moreover, an average direct effect (ADE; the effect of repetition number on parietal alpha ERD after considering pre-stimulus parietal alpha power) was significant in young subjects (β = 0.17; p = 2e-16; CI = [0.13, 0.21]) and older subjects (β = 0.04; p = 0.002; CI = [0.01, 0.06]) (Supplementary Figure 5).

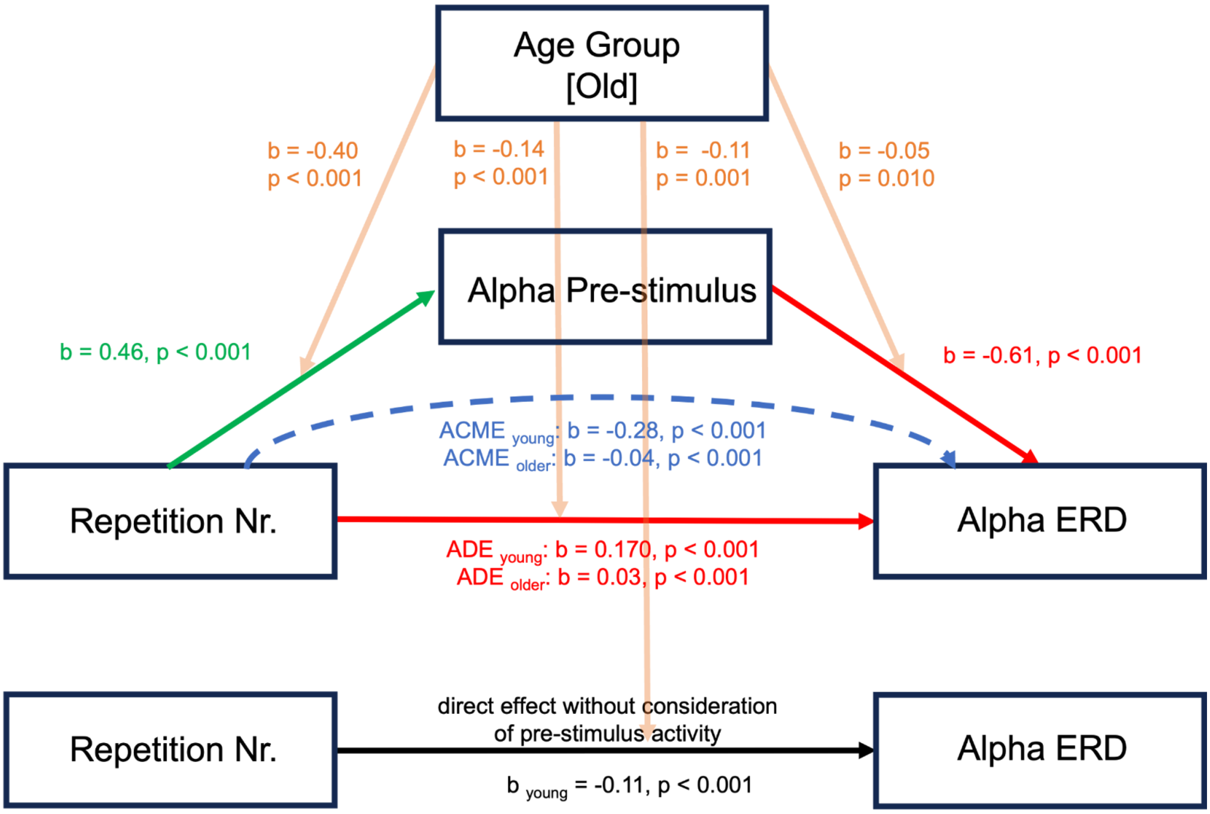


Supplementary Figure 5. Moderated mediation analysis investigating the effects of pre-stimulus parietal alpha power on parietal alpha ERD. The results show that a part of the total effect of repetition number on parietal alpha ERD is mediated by pre-stimulus parietal alpha power.

#### 5. 7. Aperiodic component

While the aperiodic component was not the primary focus of this work, we provide a brief description of the results here for completeness. The detailed analysis and interpretation of the aperiodic signal remain outside the scope of this paper and are not included in the main text. Using the sliding window approach, we obtained aperiodic component estimates (i.e., aperiodic offset and exponent) ranging from -500 to 900 ms around stimulus onset, in 50 ms intervals, for each channel and sequence repetition. We next averaged the estimates over parieto-occipital electrode clusters (i.e., identical to parietal alpha electrodes) and plotted the progression of aperiodic offset and exponent over time and sequence repetitions (Supplementary Figure 6). The plot revealed very similar progressions of aperiodic offset and exponent, characterized by event-related increase of both signals shortly after stimulus offset. Based on visual inspection, we fitted linear mixed-effects models to investigate pre-stimulus (-500 to -250 ms), event-related increase (0 to 500 ms), and late (500 to 900 ms) modulations of the aperiodic exponent over sequence repetitions and age groups (Supplementary Tables 11, 12 and 13). The models revealed that older subjects exhibit flatter exponents than their younger counterparts in all three time windows. Moreover, the models indicated a significant positive effect of repetition number on the aperiodic exponent in both the pre-stimulus and late time windows, suggesting an increasing aperiodic exponent over the course of learning. However, in both cases, the effect was weaker in older subjects, as indicated by a significant interaction between repetition number and age group.

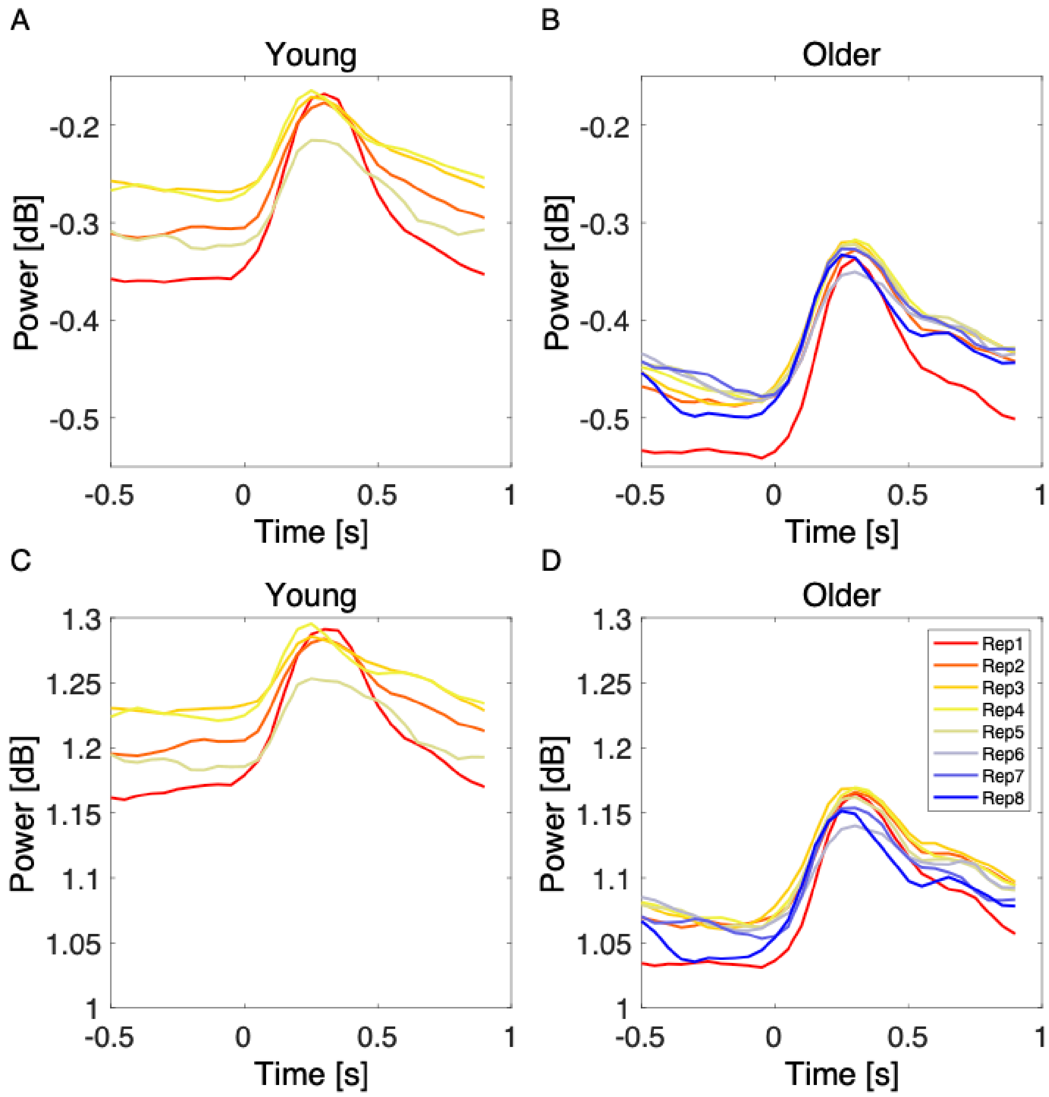


Supplementary Figure 6. Aperiodic offset (A-B) and exponent (B-C) over sequence repetitions in young and older subjects. On average, older subjects exhibited lower offset and flatter exponent compared to younger counterparts.

Supplementary Table 11. Effects of repetition number and age group on pre-stimulus aperiodic exponent

| ***Variable*** | ***β*** | ***SE*** | ***CI*** | ***t-value*** | ***p-value*** |
|---|---|---|---|---|---|
| Intercept | -0.18 | 0.05 | -0.27 – -0.08 | -3.61 | 2.9e-4*** |
| RepetitionNr | 0.03 | 0.01 | 0.02 – 0.04 | 5.91 | 2.53e-9*** |
| AgeGroup [Old] | -0.18 | 0.03 | -0.24 – -0.12 | -5.83 | 5.98e-9*** |
| RepetitionNr*AgeGroup [Old] | -0.01 | 0.01 | -0.01 – -0.00 | -2.68 | 0.007** |

| Variance components | SD | Goodness of fit | |
|---|---|---|---|
| Subject | 0.22 | Log likelihood | 1591.6 |
| Residual | 0.17 | | |

*Note. Note. Intercept represents the first repetition of young participants. RepetitionNr = Repetition number. β = unstandardized regression coefficient. SE = Standard error. CI = Confidence interval. SD = Standard deviation.*

Supplementary Table 12. Effects of repetition number and age group on event-related aperiodic exponent

| ***Variable*** | ***β*** | ***SE*** | ***CI*** | ***t-value*** | ***p-value*** |
|---|---|---|---|---|---|
| Intercept | -0.05 | 0.05 | -0.14 – 0.05 | -1.04 | 0.298 |
| RepetitionNr | 0.00 | 0.00 | 0.00 – 0.02 | 1.21 | 0.225 |
| AgeGroup [Old] | -0.19 | 0.03 | -0.25 – -0.13 | -6.05 | 1.57e-9*** |
| RepetitionNr*AgeGroup [Old] | -0.00 | 0.00 | -0.00 – 0.01 | 0.38 | 0.701 |
| **Variance components** | **SD** | **Goodness of fit** | | | |
| Subject | 0.22 | Log likelihood | 2254.8 | | |
| Residual | 0.15 | | | | |

*Note. Note. Intercept represents the first repetition of young participants. RepetitionNr = Repetition number. β = unstandardized regression coefficient. SE = Standard error. CI = Confidence interval. SD = Standard deviation.*

Supplementary Table 13. Effects of repetition number and age group on late aperiodic exponent

| ***Variable*** | ***β*** | ***SE*** | ***CI*** | ***t-value*** | ***p-value*** |
|---|---|---|---|---|---|
| Intercept | -0.16 | 0.5 | -0.26 – -0.06 | -3.19 | 0.001** |
| RepetitionNr | 0.02 | 0.00 | 0.01 – 0.03 | 5.59 | 2.41e-8*** |
| AgeGroup [Old] | -0.16 | 0.03 | -0.22 – -0.10 | -5.00 | 5.76e-7*** |
| RepetitionNr*AgeGroup [Old] | -0.01 | 0.00 | -0.01 – -0.00 | -2.80 | 0.005** |
| **Variance components** | **SD** | **Goodness of fit** | | | |
| Subject | 0.22 | Log likelihood | 1912.2 | | |
| Residual | 0.16 | | | | |

*Note. Note. Intercept represents the first repetition of young participants. RepetitionNr = Repetition number. β = unstandardized regression coefficient.  SE = Standard error. CI = Confidence interval. SD = Standard deviation.*
